# Analysis of sex differential genes at whole genome level reveals their organ-and chromosome-dependent expressions, regulations, and interactions

**DOI:** 10.1101/2022.09.24.509319

**Authors:** Jiamin Ma, Lishi Wang, Gang Yuan, Chrishan Fernando, Lan Yao, Wei Dong, Cheng Tian, Anna Bukiya, Monica M. Jablonski, Helin Feng, Daniel Z. Li, Lu Lu, Yan Jiao, Jiafu Ji, Gang Wang, Weikuan Gu

## Abstract

Males and females possess genomes that are almost identical but differ in morbidity, prevalence, severity, response to therapies and mortality of many diseases. We comprehensively analyzed gene expression data in seven tissues from BXD recombinant inbred (RI) mouse strains. We found that there were considerable differences in the numbers and functions among different tissues and between autosomal and sex chromosomes. Among sex differential genes, those on autosomal chromosomes mainly function to regulate metabolic pathways related to their host organs, while those on sex chromosomes mainly regulate nuclear proteins required for DNA replication or transcription during early development. Kidney possessed the fewest sex differential genes on sex chromosomes. The sexually dimorphic genes expressed on X chromosomes were all related to perception, while genes on the autosomal chromosomes were related to metabolism. These patterns of gene expression do not translate into similarity of protein structures, rather they are grouping with function.

## 1 Introduction

Sex differences relate to many human activities and diseases. One of the most obvious sex related differences include susceptibility to disease. In some cases, unique anatomical differences lead to women having a higher risk of breast cancer than men while they have no risk of prostate cancer, which occurs only in men (Siegel et al., 2017; Torre et al., 2015). Even in organ systems that men and women share, sex-specific differences in susceptibility to disease remain. For example, the third most diagnosed cancer among men and women in the U.S. (Siegel et al., 2022) and worldwide is colorectal cancer, which exhibits sex- and gender-specific differences (Gabriele et al., 2016). In 2022, there were projected to be 151,030 new individual colorectal cancer diagnoses and 52,580 deaths, where 53% of new cases and 54% of deaths would occur in men, compared with 47% and 46%, respectively, predicted to occur in women (Siegel *et al*., 2022). Even without considering lifestyle behaviors, ethnicity, or regional differences, men and women differ in tumor location, morbidity, response to therapies, and mortality. Diseases that affect immune responses such as arthritis are also influenced by sex differences, (Kim and Kim, 2020). Therefore, a better understanding of biological differences between men and women in their responses to disease might lead to the development of improved targeted therapies (Gabriele *et al*., 2016).

Physical strength and activities are particularly influenced by sex differences. Learning abilities such as female verbal advantage and male navigation strategy are also sex dependent (Boone et al., 2018; Scheuringer et al., 2017). However, there remains an incomplete understanding of the systematic molecular network that exists between specific pathways that exhibit differences between sexes and their relation to hormones, particularly in the tissue-specific profiles of gene expression in different activities and diseases that are sex-specific.

Females and males often differ extensively in their physiological characteristics and physical functions (Parsch and Ellegren, 2013). For example, age-related diseases and life span exhibit noticeable sex biases, immunological responses to foreign and self-antigens show distinctions based on sex, and insulin secretion and metabolism associate with gene-specific sex differences (Ashbrook et al., 2021; Hall et al., 2014; Klein and Flanagan, 2016). This sexual dimorphism is primarily caused by differences in sex-biased gene expression, yet males and females share much of their genome (Ingleby et al., 2015; Parsch and Ellegren, 2013).

More evidence has revealed the features and degree of sex-biased gene expression in a wide range of given diverse species. The present GTEx (genotype-tissue expression) project has been devoted to characterizing variation in gene expression levels across diverse tissues of the human body and sex was one of the important factors that was considered (GTEx Consortium, 2017; LopesRamos et al., 2020). Genes show highly differential expression across individuals and low variance across tissues including autosomal genes as well as genes on the sex chromosomes (Care et al., 2018; Voskuhl et al., 2018). Specificity of tissue is likely to be driven by the collaborative expression of multiple genes in multiple cell types (Mele et al., 2015). However, due to the heterozygosity of the human population and the relatively low abundance of specific gene variants, it is often difficult to identify the causal variants and define their genetic risk across populations (Gay et al., 2020).

Rodent models such as the recombinant inbred (RI) strains derived from C57BL/6J and BA/2J (BXD) mice have contributed tremendously to our understanding of human genetics and genomics. The BXD family comprises the largest RI mouse population and offers remarkable data derived from whole genome expression profiles and phenotypes (Cao et al., 2018). The existence of tissuespecific sex-biased expression in mice means that this wealth of data is directly relevant to the human genome and provides a starting point for identifying the analogous human genes. (Crowley et al., 2015; Werner et al., 2017). Therefore, we comprehensively analyzed the data of gene expression studies in seven tissues (adrenal gland, amygdala, hypothalamus, pituitary gland, kidney, spleen, and liver) from male and female mice from the BXD family. By analyzing available RNAsequencing (RNA-seq) data from 97 strains of male and female mice, we found thousands of genes differentially expressed in these tissues that are common to both sexes. In this study, we report the analytic results of sexually dimorphic expression of genes (DEGs) in these BXD strains and describe confirmation data from human samples.

## 2. Results

The gene expression data in GeneNetwork version 1 was generated using a total of 206,523 probes that detect the genes across all the autosomes (chromosomes 1∼19) and the X chromosome, with 198,291 probes complementary to autosomes and 8,232 probes complementary to the X chromosome (Mulligan et al., 2017). The gene expression data for seven tissues common to males and females was grouped by sex. We extracted the probes whose false discovery rate (FDR) of gene expression between female and male mice was less than 0.05 and identified 3,846 autosomal probes and 767 X chromosome probes that accounted for 1.94% and 9.32% of sequences on the autosomes and X chromosome, respectively (see Table 1). Among these tissues we found more than 100 genes with extremely low p values.

**Table 1.** Number of probes that q value (FDR-adjusted p value) of gene expression is less than 0.05

| Tissue | Adrenal | Amygdala | Hypothalamus | Kidney | Liver | Pituitary | Spleen |
| --- | --- | --- | --- | --- | --- | --- | --- |
| n/N<sup>a</sup>(%) |  |  |  |  |  |  |  |
| **ChrX** | 284/1207(23.5<br>3) | 10/1207(0.8<br>3) | 10/1207(0.83<br>) | 3/1442(0.21) | 198/691(28.65) | 249/1207(20.6<br>3) | 13/1271(1.02<br>) |
| **Chr1-19** | 67/27031(0.25<br>) | 0/27031(0) | 0/27031(0) | 86/42996(0.2<br>0) | 3648/18040(20.2<br>2) | 19/27031(0.07<br>) | 26/28588(0.0<br>9) |
| **Chr1** | 2/1738(0.12) | 0/1738(0) | 0/1738(0) | 2/2278(0.07) | 239/1110(21.53) | 3/1738(0.17) | 5/1760(0.28) |
| **Chr2** | 5/2274(0.22) | 0/2274(0) | 0/2274(0) | 1/3592(0.03) | 279/1598(17.46) | 2/2274(0.09) | 0/2334(0) |
| **Chr3** | 1/1343(0.07) | 0/1343(0) | 0/1343(0) | 11/2306(0.48<br>) | 193/912(21.16) | 1/1343(0.07) | 1/1390(0.07) |
| **Chr4** | 4/1753(0.23) | 0/1753(0) | 0/1753(0) | 7/2715(0.26) | 221/1115(19.82) | 1/1753(0.06) | 1/1810(0.06) |
| **Chr5** | 5/1634(0.31) | 0/1634(0) | 0/1634(0) | 6/2826(0.21) | 228/1159(19.67) | 3/1634(0.18) | 1/1682(0.06) |
| **Chr6** | 28/1520(0.59) | 0/1520(0) | 0/1520(0) | 2/2501(0.08) | 225/1084(20.76) | 1/1520(0.07) | 4/1553(0.06) |
| **Chr7** | 4/2430(0.16) | 0/2430(0) | 0/2430(0) | 1/3001(0.03) | 323/1554(20.79) | 0/2430(0) | 0/2576(0) |
| **Chr8** | 0/1382(0) | 0/1382(0) | 0/1382(0) | 4/2214(0.18) | 174/924(18.83) | 0/1382(0) | 0/1398(0) |
| **Chr9** | 3/1595(0.19) | 0/1595(0) | 0/1595(0) | 2/2403(0.08) | 215/1060(20.28) | 22/1595(0.13) | 0/1607(0) |
| **Chr10** | 4/1299(0.31) | 0/1299(0) | 0/1299(0) | 2/2071(0.10) | 174/847(20.54) | 0/1299(0) | 0/1321(0) |
| **Chr11** | 0/2020(0) | 0/2020(0) | 0/2020(0) | 6/3421(0.18) | 270/1447(18.66) | 2/2020(0.10) | /2053(0.05) |
| **Chr12** | 0/995(0) | 0/995(0) | 0/995(0) | 2/1699(0.12) | 122/618(19.74) | 0/995(0) | 2/1019(0.20) |
| **Chr13** | 5/1076(0.46) | 0/1076(0) | 0/1076(0) | 8/1799(0.44) | 157/684(22.95) | 2/1076(0.19) | 4/1107(0.36) |
| **Chr14** | 0/1122(0) | 0/1122(0) | 0/1122(0) | 0/1648(0) | 136/653(20.83) | 0/1122(0) | 5/1175(0.43) |
| **Chr15** | 0/1061(0) | 0/1061(0) | 0/1061(0) | 11/1770(0.62<br>) | 158/713(22.16) | 1/1061(0.09) | 1/1067(0.05) |
| **Chr16** | 3/904(0.33) | 0/904(0) | 0/904(0) | 16/1539(1.04<br>) | 120/614(19.54) | 0/904(0) | 0/907(0) |
| **Chr17** | 2/1305(0.15) | 0/1305(0) | 0/1305(0) | 0/1924(0) | 185/853(21.69) | 0/1305(0) | 0/1323(0) |
| **Chr18** | 6/673(0.89) | 0/673(0) | 0/673(0) | 0/1271(0) | 74/471(15.71) | 0/673(0) | 4/682(0.58) |
| **Chr19** | 14/907(1.54) | 0/907(0) | 0/907(0) | 5/1520(0.33) | 155/624(24.84) | 1/907(0.11) | 0/914(0) |

a. N-the total number of probes; n-the number of probes which the gene expression's q value (FDR-adjusted p value) is less than 0.05.

### 2.1 Differential expression of genes on autosomal chromosomes across tissues

The number of probes that represent the differential expression (p value less than 0.05) across each autosome in seven tissues is shown in Table 1. Interestingly, we identified no differential expression between the sexes in autosomes in the amygdala and hypothalamus, while approximately 30% of the probes (5,897 of 19,493) specific for liver tissue exhibited gene expression that varied in the two sexes, corresponding to 5,407 identifiable unique genes.

The proportion of the probes specific for other tissues that detected differentially expressed genes (DEGs) ranged from low to moderate. For example, ∼2% (547 of 27,031) of probes detected 513 identifiable unique DEGs in the adrenal gland, ∼1% (415 of 42,996) of probes detected 325 identifiable unique DEGs in kidney, ∼1% (268 of 27,031) of probes detected 191 identifiable unique DEGs in pituitary gland, and 7.3% (2,023 of 27,678) of probes detected 1,883 identifiable unique DEGs in spleen.

The DEGs whose *p* values were so low that they approached zero for expression in the adrenal gland were the *Akr1d1* (aldo-keto reductase family 1 member D1; also known as steroid A-ring 5βreductase) gene on mouse chromosome (Chr 6) and the *Akr1c18* (aldo-keto reductase family 1, member C18) gene on Chr 13; for expression in the kidney were *Slc7a12* (solute carrier family 7 (cationic amino acid transporter, y+ system), member 12) and *Cyp7b1* (cytochrome P450 family 7 subfamily B member 1) genes on Chr 3, the *Slc7a13* gene on Chr 4, *Slco1a1* (solute carrier organic anion transported family member 1A1) on Chr 6, and *Acsm3* (acetyl-CoA synthetase medium chain family member 3) on Chr 7; for expression in the pituitary gland, *Rxfp1* (relaxin family peptide receptor 1) on Chr 3; and for expression in liver, there were 119 identifiable genes. All the false discovery rate (FDR) q values (FDR-adjusted *p*-values) were less than 0.05. The protein products from these genes represent no more than 11.7 % of amino acid identity and no more than 28.9% of amino acid similarity (**Supplementary Figure D**), both of which are well below the required 40% of sequence identity for predicting similar functionality (Pearson, 2013).

#### 2.1.1 *Akr1d1* is the most different expressed genes between male and female in adrenal glands

*Akr1d1* and *Akr1c18* were differentially expressed (*p* = 0) in adrenal glands from male and female mice. Because humans lack the *Akr1c18* gene, we focused on *Akr1d1* for detailed analysis. A total of 18 datasets of *Akr1d1* expression levels were obtained from seven tissues (liver and kidney have distal and mid 3’ UTR) (**Supplementary table 1**). The mean expression levels in seven tissues from each sex showed variation in the magnitude of the difference (Figure 1A). While the sex difference in three tissues (liver, kidney, and adrenal glands) reached statistical significance, the differences observed in the adrenal gland were the largest (Figure 1B). When we calculated Pearson’s correlation coefficient of *Akr1d1* expression in all seven of the tissues we examined, its expression levels in kidney showed high positive correlation with those in liver and adrenal glands and between male and female mice within the kidney (Figure 1C). In contract, there was no correlation between its expression levels in the adrenal gland from female and male mice (Figure 1D). In adrenal gland, the top 100 most correlated phenotypes to *Akr1d1* in male mice were weight loss (WL) with a positive correlation (r = 0.890), while the top correlated phenotypes in female mice were sibling sucking (SIB) with a negative correlation (r = −0.948); WL exhibited a non-significant positive correlation (r = 0.239) to SIB. Among the top 100 most correlated genes to *Akr1d1* in female mice, *Atp8b1* (ATPase phospholipid transporting 8B1; r = 0.843) exhibited a non-significant negative opposite (r = −0.219) correlation to that of *Cyp21a1* (cytochrome P450 family 21, subfamily a, polypeptide 1; r = 0.947) in male mice and *vice versa* (Figure 1E). To further understand the relationship of *Akr1d1* expression in male versus female mice, we examined the top genotypes and genes from the adrenal gland and the other six tissues (Figure 1F). The results showed that the strong correlation between *Akr1d1* and its top corelated genes and phenotypes in adrenal gland did not hold for other tissues, where the correlation varied greatly. While these correlations were much less in all other tissues, the correlations observed in kidney were relatively similar to those in the adrenal gland, except that the SIB phenotype in female mice exhibited positive correlation to the kidney but negative correlation to the adrenal gland.

**Figure 1.**
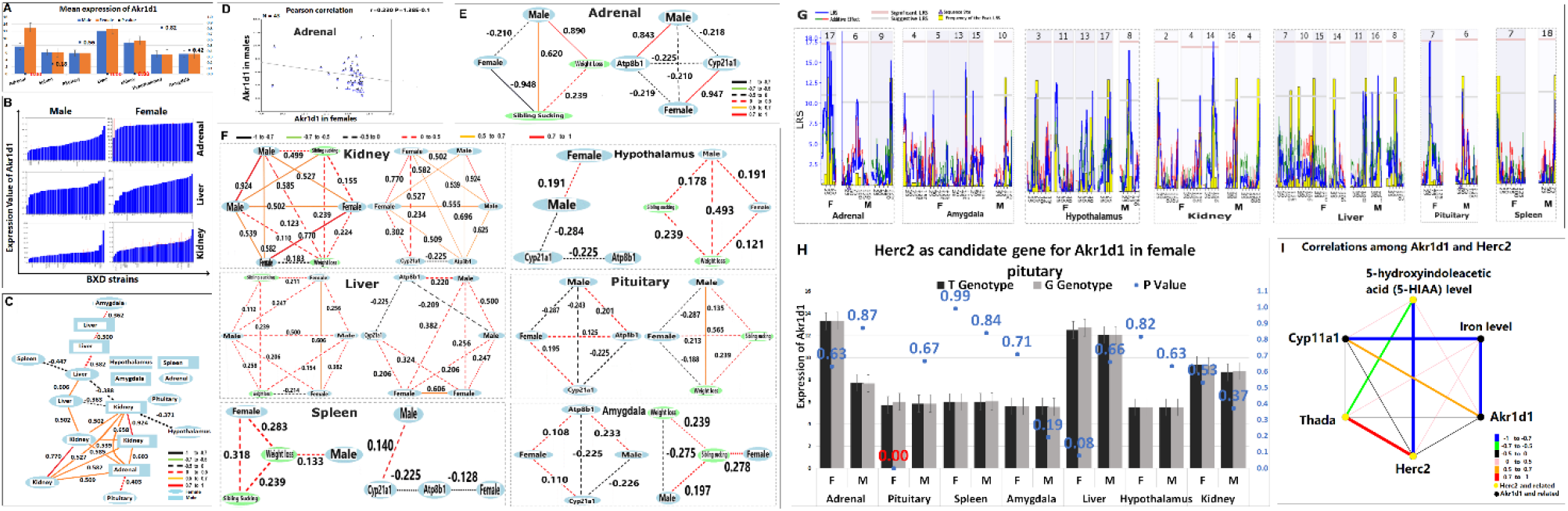
Expression of *Akr1d1*. **A.** Mean expression of *Akr1d1* in all tissues examined: adrenal glands, spleen, pituitary gland, liver, kidney, hypothalamus, and amygdala. The blue and tangerine clustered columns represent the mean expression values for *Akr1d1* in male (blue) and female (tangerine) mice, respectively. The y-axis on the left represents the relative expression of *Akr1d1*, while that on the right represents the *p* values. The scatter markers represent the *p* values calculated by the student’s t-test; statistically significant differences (*p* < 0.05) are shown in red. **B.** Differential *Akr1d1* expression in liver, kidney, and adrenal glands of male and female BXD mice. **C.** The networks of *Akr1d1* in all tissues. *Akr1d1* expression in kidney shows significant positive relationship with that in liver and adrenal gland. Within kidney, the expression levels of *Akr1d1* in males and females were also positively correlated. **D.** The Pearson correlation of *Akr1d1* in adrenal glands between male and female mice. **E.** The correlations between *Akr1d1* and its most related genes and phenotypes in adrenal glands. The most correlated phenotype to *Akr1d1* in male mice was weight loss (WL) with r = 0.89, while that in female mice was sibling sucking (SIB) with r = −0.948; WL exhibited a slight positive correlation (r = 0.239) to SIB. The most correlated genes to *Akr1d1* in female mice was *Atp8b1* (r = 0.843) and in male mice was *Cyp21a1*(r = 0.947), both of these genes show a slight opposite (r = −0.219) correlation between them. **F.** The networks among *Akr1d1*, WL, SIB, *Atp8b1*, and *Cyp21a1* in other tissues. In the kidney of male mice, *Akr1d1* was positively correlated to SIB. *Atp8b1* showed positive correlations between male and female mice. Similar to the pattern observed in the adrenal gland, expression of *Akr1d1* in pituitary gland from male mice was positively correlated to WL. We detected no strong correlations between the sexes in hypothalamus, liver, spleen, and amygdala. **G.** The e-QTL maps of *Akr1d1* in all tissues. Only the chromosomes that exhibited LRS levels higher than “suggestive” or “significant” were shown in the figure. Red and green lines represent the additive genetic contribution; red lines indicate negative values (B6 alleles with increasing trait values) and green lines indicate positive values (D2 alleles with increasing trait values). The yellow bars represent the relative frequency of peak LRS at a given location from 2,000 bootstrap resamples, the gray bars represent the suggestive level, and the pink bars represent the significance level. The orange hash marks on the x-axis signify the SNP density (sequence difference between the two parental strains). The purple triangles on the x-axis represent the location of *Akr1d1*. Three chromosomes exhibited e-QTLs higher than significant LRS level: Chr 17 of female adrenal gland, Chr 9 of female kidney, and Chr 6 of female pituitary gland. **H.** The expression value of *Akr1d1* in C57BL/6J (B6; T) and DBA/2J (D2; G) genotypes. The student’s t-test was used to calculate the difference in expression values between these genotypes; statistically significant differences (*p* < 0.05) are shown in red. Judging by its *p* value, *Herc2* is a candidate regulator gene for *Akr1d1* in the adrenal gland of female mice. **I.** The correlations between *Akr1d1* and *Herc2*, including their most related genes and phenotypes. The most related genes and phenotypes to *Akr1d1* in the adrenal gland were *Cyp11a1* and 5hydroxyindoleacetic acid (5-HIAA) levels, respectively. The most related genes and phenotypes to *Herc2* in the adrenal gland were *Thada* and *Herc2*. We detected no strong correlations between *Akr1d1* and *Herc2*.

To further understand the relationships between phenotypes and genotypes, we determined the expression quantitative trait loci (e-QTL) of *Akr1d1* in adrenal gland and other tissues and mapped the loci whose likelihood ratio statistics (LRS) levels were higher than the “suggestive” threshold level (Figure 1G). The results of the e-QTL analysis suggested that the expression of *Akr1d1* from Chr 6 might be modulated in female mice from a distal region on Chr 17 and in male mice from local regions on Chr 6 and distal region on Chr 9. Mapping of the *Akr1d1* e-QTL in the other tissues from female mice (Figure 1G) located eQTLs on three chromosomes (Chr 17 in the adrenal gland, Chr 14 in the kidney, and Chr 7 in the pituitary gland), with LRS levels higher than the “significant” threshold level (**Supplementary figure A-C**). Using conducting interval and haplotype analysis of these tissues from female mice, we identified nine SNPs and 339 genetic elements on Chr 17 of the adrenal gland (**Supplementary table 2-3**), five SNPs but no genes on Chr 14 of the kidney (**Supplementary table 4**), and six SNPs and 16 genes on Chr 7 of the pituitary gland (**Supplementary table 5-6**). Among the above genetic elements, only two of which contained SNPs: *Cchcr1* (coiled coil alpha-helical rod protein 1) contained SNP rs13482963 and *Herc2* (HECT and RLD domain containing E3 ubiquitin protein ligase 2) contained SNP rs32328305.

The eQTL maps (**Supplementary figure A, C**) localized SNP rs32328305 in the mouse *Herc2* gene to the “peak” LRS locus, which predicts a high probability that SNP rs32328305 regulates the expression of *Akr1d1*. The rs32328305 SNP is a T-to-G substitution in the intron region discovered in B6 (T) and D2 (G) mouse strains. Introns as non-coding region are expected to possibly contribute to chromatin structures, transcription regulation, RNA splicing and the translational machinery (Nakagawa and Fujita, 2018). When we compared the expression of the T and G *Akr1d1* genotypes, only the pituitary gland of female mice exhibited a significant difference in expression levels (*p* = 0.0002), suggesting that the *Herc2* gene or its protein product may regulate the expression of *Akr1d1* in the pituitary gland of female mice (Figure 1H).

The top related phenotype of *Akr1d1* in the pituitary gland of female mice was the iron level (IL) in the medial prefrontal cortex and the top related genotype was *Cyp11a1* (cytochrome P450 family 11 subfamily A member 1). The top related phenotype of *Herc2* was the level of 5hydroxyindoleacetic acid (5-HIAA) in the blood of adult female mice and the top related genotype of *Herc2* was *Thada* (THADA armadillo-repeat containing gene). Mapping of the networks among *Akr1d1* and *Herc2*, including their top related genes and phenotypes showed no strong correlations between *Akr1d1* and *Herc2* (Figure 1I).

#### 2.1.2 The differentially expressed genes in kidney are in similar locations and have strong correlations

We found that five DEGs were differentially expressed (*p* = 0) in the kidneys of male and female mice: *Slc7a12* and *Cyp7b1* on Chr 3, *Slc7a13* on Chr 4, *Slco1a1* (ID: 1420379_at) on Chr 6, and *Acsm3* on Chr 7. There is strong correlation among *Cyp7b1*, *Slc7a13*, and *Slco1a1* (Figure 2A) and their most frequent phenotypes exhibit similarities (Table 2). According to Andrew Ransick (Ransick et al., 2019) *Cyp7b1*, *Slc7a13,* and *Slc7a12* are localized to the kidney proximal tubule clusters where the expression levels of several other genes exhibit sex bias in mice. As *Slc7a12* does not exist in *Homo sapiens*, we will not consider it further here.

**Figure 2.**
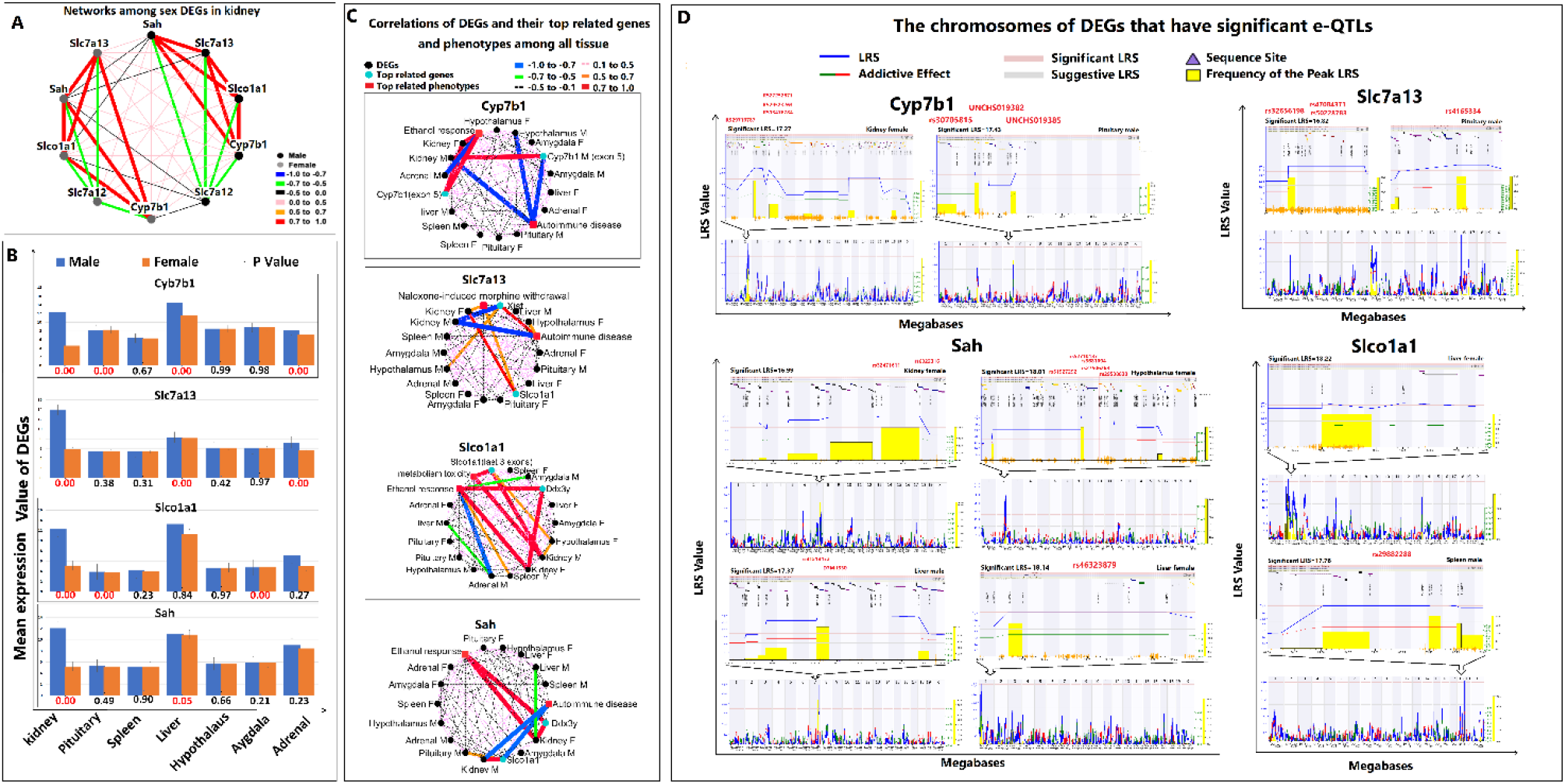
Sexually dimorphic gene expression in mouse kidney. **A.** The networks of kidney DEGs. The black circles (nodes) represent the genes that were expressed at higher levels in male mice, the gray circles (nodes) represent the genes that were expressed at higher levels in female mice, and the edge (line) that connects the nodes represents the sample correlation, rho (r). We discovered strong correlations among the DEGs in male and female mice. **B.** Mean expression values of kidney DEGs in all tissues examined. The panel is labeled as described in the legend to Figure 1A, except that the y-axis represents the mean expression values for each DEG. *Cyp7b1* exhibited sex bias in kidney, pituitary gland, liver, and adrenal gland; *Slc7a12* and *Slco1a1* exhibited sex bias in kidney, liver, and adrenal gland; and *Acsm3* exhibited sex bias in kidney. **C.** The networks of kidney DEGs in all tissues. The black circles represent the DEGs, the blue circles represent their most related genes, the red rectangles represent their most related phenotypes, and the edge that connects the nodes represents the sample correlation. From the top, the first network shows the correlations of *Cyp7b1*. The top related phenotypes of *Cyp7b1* in kidney were the genes that expressed on exon 5. The top related phenotypes of *Cyp7b1* in the kidney of male mice was autoimmune disease (AD), while in the kidney of female mice was ethanol response (ER). In male mice, AD had strong negative correlation to *Cyp7b1* in the hypothalamus and ER had strong negative correlation to *Cyp7b1* in the adrenal gland. The second network from the top shows the correlations of *Slc7a13*. *Xist* and *Slco1a1* were the most related genes to *Slc7a13* in the kidney of male and female mice, respectively. Autoimmune disease (AD) and morphine withdrawal (MW) were the most related phenotypes to *Slc7a13* in the kidney of male and female mice, respectively. In the hypothalamus, *Xist* had positive correlation to *Slc7a13* in male mice, while AD had positive correlation to *Slc7a13* in female mice. The network third from the top shows the correlations of *Slco1a1*. The top related genes to *Slco1a1* in kidney are the last three exons of *Slco1a1* in male mice and *Ddx3y* in female mice. The last three exons of *Slco1a1* had positive correlations to the hypothalamus female of mice. In the kidney, the top related phenotypes of *Slco1a1* were metabolism toxicity (MT) in male mice and ethanol response (ER) in female mice. In male mice, ER had a negative correlation to *Slco1a1* in the adrenal gland and positive correlations to *Slco1a1* in the spleen. The bottom network shows the correlations of *Acsm3*. In kidney, the top related genes to *Acsm3* were *Slco1a1* in male mice and *Ddx3y* in female mice. Autoimmune disease (AD) was the top related phenotype of *Slco1a1* in the kidney of male mice, while ethanol response (ER) was the top related phenotypes of *Slco1a1* in the kidney of female mice. **D.** The e-QTL maps of DEGs. For each panel, the interval map of the sequence in the entire mouse genome is shown at bottom, while the map within a significant QTL location is shown at the top. The figure is labeled as described in the legend to Figure 1G, except that the SNPs in the maximal LRS location and those that occur in genes are labeled in red and the red vertical lines indicate whether the SNPs co-localize with the frequency of peak LRS location. The *p* value for each gene is listed on the top left of each map and the tissue information is provided at top right. The upper left panel shows the e-QTL maps of *Cyp7b1* in the kidney of female mice (left) and in the pituitary gland of male mice (right). UNCHS019385 did not localize to the peak LRS location. The upper right panel shows the e-QTL maps of *Slc7a13* in the pituitary gland of male mice. Only rs4165334 localizes to the peak LRS location. The lower left panel shows the e-QTL maps of *Acsm3* in the kidney and liver of male mice and the hypothalamus and liver of female mice. In female kidney (upper left), both SNPs localized to the peak LRS location. In female hypothalamus (upper right), only rs29538830 did not localize to the peak LRS location. In the liver, both SNPs detected in male mice (bottom left) localized to the peak LRS location, whereas the only SNP detected in female mice (bottom right) did not localize to the peak LRS location.

**Table 2:**
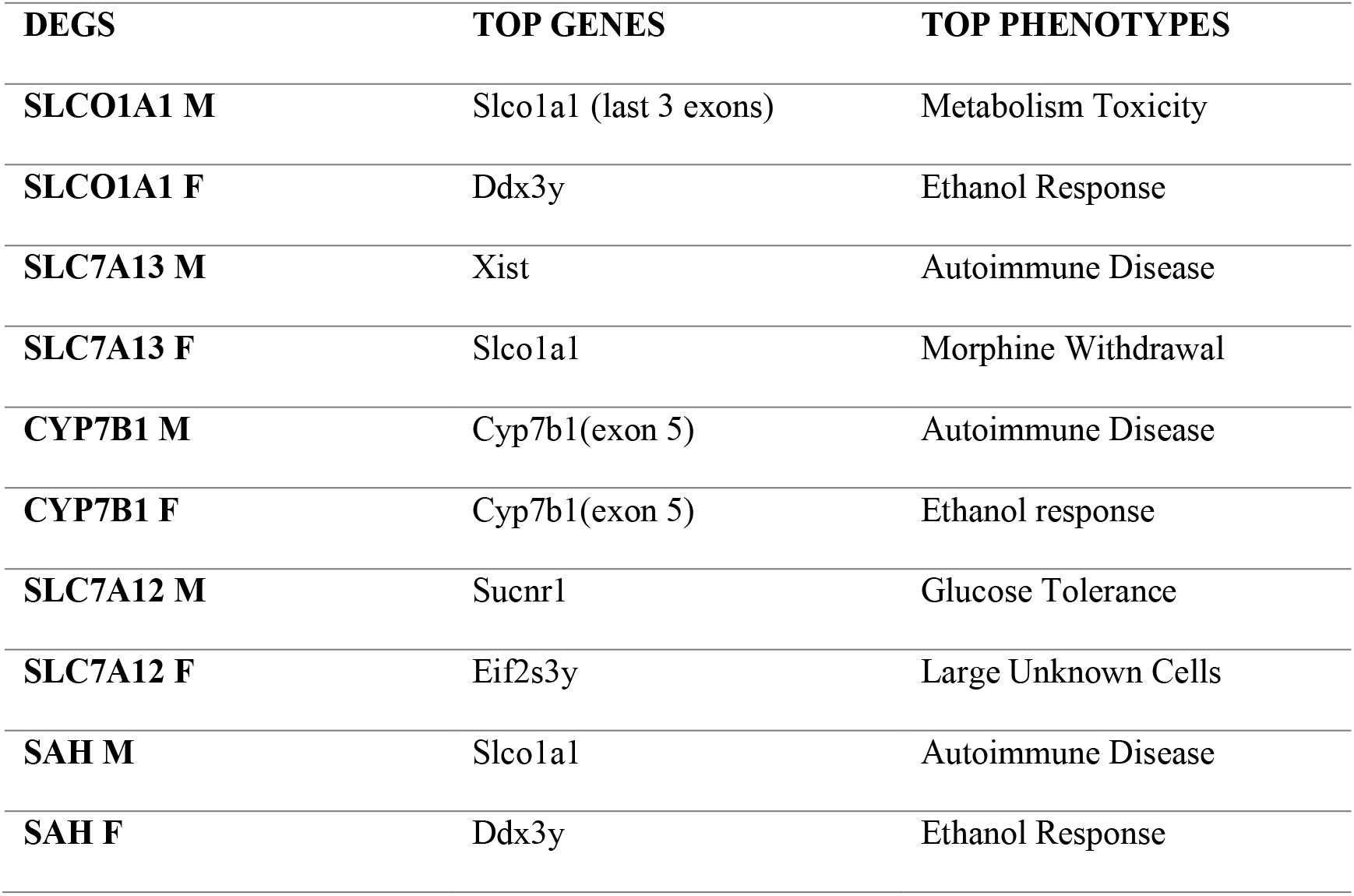
The stop related gens and phenotypes of sex DEGs in kidney

| DEGS | TOP GENES | TOP PHENOTYPES |
| --- | --- | --- |
| SLCO1A1 M | Slco1a1 (last 3 exons) | Metabolism Toxicity |
| SLCO1A1 F | Ddx3y | Ethanol Response |
| SLC7A13 M | Xist | Autoimmune Disease |
| SLC7A13 F | Slco1a1 | Morphine Withdrawal |
| CYP7B1 M | Cyp7b1(exon 5) | Autoimmune Disease |
| CYP7B1 F | Cyp7b1(exon 5) | Ethanol response |
| SLC7A12 M | Sucnr1 | Glucose Tolerance |
| SLC7A12 F | Eif2s3y | Large Unknown Cells |
| SAH M | Slco1a1 | Autoimmune Disease |
| SAH F | Ddx3y | Ethanol Response |

The expression of *Cyp7b1* (ID: 1421074_at), *Slc7a13* (ID: 1449301_at), *Slco1a1* (ID: 1420379_at) and *Sah* (*Acsm3)* in seven organs from male and female mice is shown in Figure 2B. In addition to kidney, *Cyp7b1* also exhibited a sex bias in pituitary gland, liver, and adrenal gland tissues, *Slc7a13* and *Slco1a1* also exhibited sex biases in liver and adrenal gland (*p* < 0.5), and *Acsm3* exhibited a sex bias in the liver.

The correlation network of top related genes and phenotypes of the DEGs in kidney and other tissues were mapped separately in Figure 2C. This network data suggested that **1)** in female mice, the phenotypes that were most positively related to *Cyp7b1*, *Slco1a1*, and *Acsm3* involved the response to ethanol. In contract, in male mice the ethanol response exhibited a strong negative correlation with solute carrier organic anion transporter family member 1D1 (*Slco1d1* in the adrenal gland (r = −0.81) and a moderate positive correlation with *Slco1d1* in the spleen (r = 0.69). **2)** in male mice, the most negatively related phenotype of *Cyp7b1*, *Slc7a13*, and *Acsm3* was autoimmune disease. While autoimmune disease exhibited a strong negative correlation with *Cyp7b1* in the hypothalamus of male mice (r = −0.83), it exhibited a moderate positive correlation with *Slc7a13* in the hypothalamus of female mice (r = 0.51). **3)** *Slco1a1* was the most related gene to itself, to *Slc7a13* from female mice, and to *Acsm3* from male mice, but in male mice was located at different locations in the genome and exhibited moderate positive correlation with *Acsm3* in the pituitary gland (r = 0.58). **4)** in female mice, *Ddx3y* (DEAD-box helicase 3, Y-linked) was the most positively related gene to *Sclo1a1* and *Acsm3* and it connected to their most positive related phenotype, ethanol response. 5) In the kidney of male mice, *Xist* (long noncoding RNA for X-chromosome inactivation) is the most negatively related gene to *Slc7a13*, which has positive correlation to *Slc7a13* in the hypothalamus of male mice (r = 0.98).

A total of 10 e-QTLs that were predicted to regulate the expression of these four DEGs from five tissues were mapped on five chromosomes (Figure 2D). *Hao1* (2-hydroxyacid oxidase 1) and *Plcb4* (phospholipase C beta 4) are candidate regulators for *Cyp7b1*. There are significant e-QTLs on Chr 2 in the kidney of female mice and on Chr 6 in the pituitary gland of male mice for predicted to regulate *Cyp7b1* expression levels. In the kidney, one SNP in *Hao1* and three SNPs in *Plcb4* were present in the maximum LRS (LRS = 22.096) and the frequency of peak LRS locations (**Supplementary table 7-8,** Figure 2D), In the pituitary gland, two SNPs occur in each *Gm6313* and *Riken* (*4930528G23Rik*) genes at both frequency of peak LRS and the maximum LRS (LRS = 18.509) locations **(Supplementary table 9-10,** Figure 2D). Some eQTLs mapped to the candidate genes *Gm6313* and *Riken*, but they were not considered further here as they do not exist in the human genome. Consequently, we hypothesized that *Hao1* and *Plcb4* were potential physiologically relevant candidate regulators for *Cyp7b1*.

When mapping e-QTLs of *Slc7a13*, we observed no significant e-QTLs in kidney but discovered that two chromosomes (Chr 8 and Chr16) exhibited significant e-QTLs in the pituitary gland from male mice (Figure 2D), where a single SNP controlled by the *Masp1* (mannan-binding lectin (MBL) serine protease 1) gene was identified as a candidate regulator for *Slc7a13* in the pituitary gland (**Supplementary table 11-12**, Figure 2D).

When mapping the e-QTLs of *Sah* (*Acsm3*), four significant e-QTLs mapped to the following locations: Chr 7 in the kidney of female mice, Chr 2 in the hypothalamus of female mice, Chr 1 and 7 in the liver of male mice. In female kidney, we identified two SNPs in the *Mosmo* (modulator of smoothened protein) and *Abca14* (ATP-binding cassette, subfamily A (ABC1), member 14) as potential candidate regulators of *Acsm3* as they both are present at the frequency of peak LRS and the maximum LRS (LRS = 23.027) locations (**Supplementary table 13**, Figure 2D). As the *Abca14* gene does not exist in the human genome, we refrained from studying it further. On Chr 2 in the hypothalamus of female mice, we discovered three SNPs occurring on two genes, solute carrier family 12 member 6 (*Slc12a6*) and the endoplasmic reticulum (ER) membrane protein complex subunit 7 precursor (*Emc7*) that were present at the frequency of peak LRS and the maximum LRS (LRS = 37.013) locations (**Supplementary table 14,** Figure 2D). In male liver, two SNPs and two genes, acyl-CoA synthetase medium chain family member 5 (Acsm5) and member 1 (Acsm1) were present at the frequency of peak LRS and the maximum LRS (LRS = 29.234) locations (**Supplementary table 15**, Figure 2D). In the liver of female mice, one SNP in one gene was present at the maximum LRS (LRS = 18.133) location but it was not found at the Peak LRS location (**Supplementary table 16**, Figure 2D).

When mapping the e-QTLs of *Slco1a1*, we also discovered that two chromosomes had significant e-QTLs: Chr 2 in the liver of female mice and Chr 18 in the spleen of male mice (Figure 2D). On Chr 2 of female liver, there is no gene was present at the maximum LRS (LRS = 19.082) location (**Supplementary table 17)**. On Chr 18 of the male spleen, there is single SNP contained two genes, *Fbxo15* (F-box protein 15) and *Timm21* (translocase of inner mitochondrial membrane 21) that were predicted to be candidate regulators of *Slco1a1* in these tissues (**Supplementary table 19)**.

Thus, we identified a total of 12 SNPs and 10 genes as candidate regulators that exist in humans (Table 3). The expression value of the DEGs between B6 and D2 genotype of candidate SNPs suggests that these candidate genes can regulate the DEGs in more than one tissue (Figure 3A). Mapping of the networks among these DEGs and the corresponding candidate regulator genes showed them to be connected by their top related genes or top related phenotypes (Figure 3B). For example, *Cyp7b1* exhibited a strong negative correlation to the top related phenotypes of *Plcb4* (liver lipidomics) and its top related phenotype, ethanol response, exhibited strong negative correlation to the top related phenotype of *Hao1*, i.e., midbrain dopamine levels (Figure 3B).

**Figure 3.**
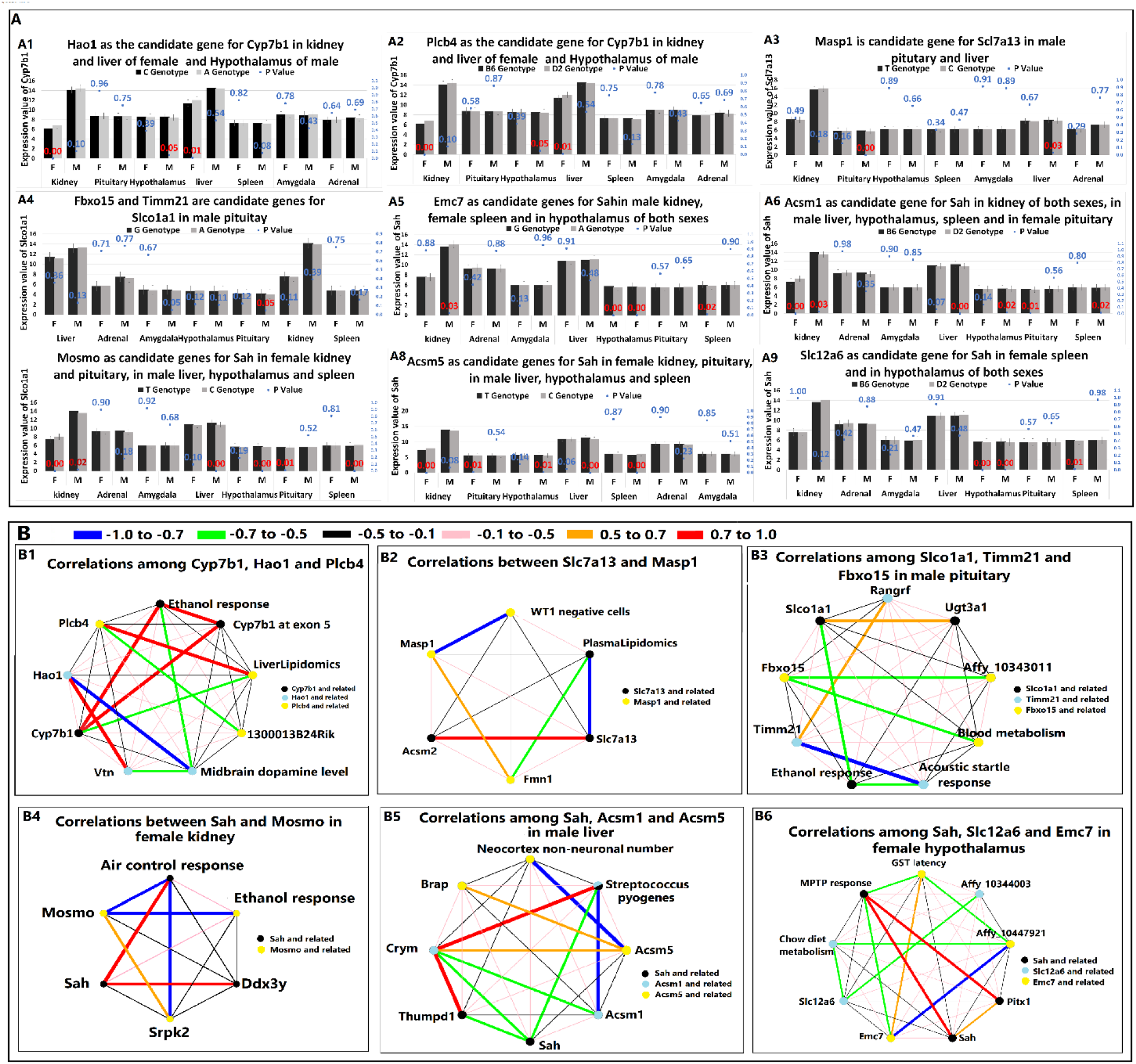
Candidate regulators of DEGs from the mouse kidney. **A.** The DEG expression of candidate regulator genes between B6 and D2 genotypes. The panel is labeled as described for Figure 1A except that the black and gray clustered columns represent the mean expression of DEGs in the B6 (black bars) and D2 (gray bars) genotypes, respectively. The title of each chart (A1-A9) indicates the tissue distribution of the candidate regulator genes: **A1,** *Hao1* candidate for *Cyp7b1*; **A2,** *Plcb4* candidate for *Cyp7b1*; **A3,** *Masp1* candidate for *Scl7a13*; **A4**, *Fbxo15* and *Timm21* candidates for *Slco1a1*; **A5**, *Emc7* candidate for *Sahin*; **A6**, *Acsm1* candidate for *Acsm3*/*Sah*; **A7**, *Mosmo* candidate for *Acsm3/Sah*; **A8**, *Acsm5* candidate for *Acsm3/Sah*; and **A9**, *Slc12a6* candidate for *Acsm3/Sah*. **B.** The correlations among DEGs and their candidate genes. The panel is labeled as described in the legend to Figure 2D. **B1**, correlations between *Cyp7b1*, *Hao1*, and *Plcb4*; **B2**, correlations between *Slc7a13* and *Masp1*; **B3**, correlations among *Slco1a1*, *Timm21*, and *Fbxo15*; **B4**, correlations between *Acsm3*/*Sah* and *Mosmo*; **B5**, correlations among *Acsm3*/*Sah*, *Acsm1*, and *Acsm5*; and **B6**, correlations among *Acsm3*/*Sah*, *Slc12a6*, and *Emc7*.

**Table 3.**
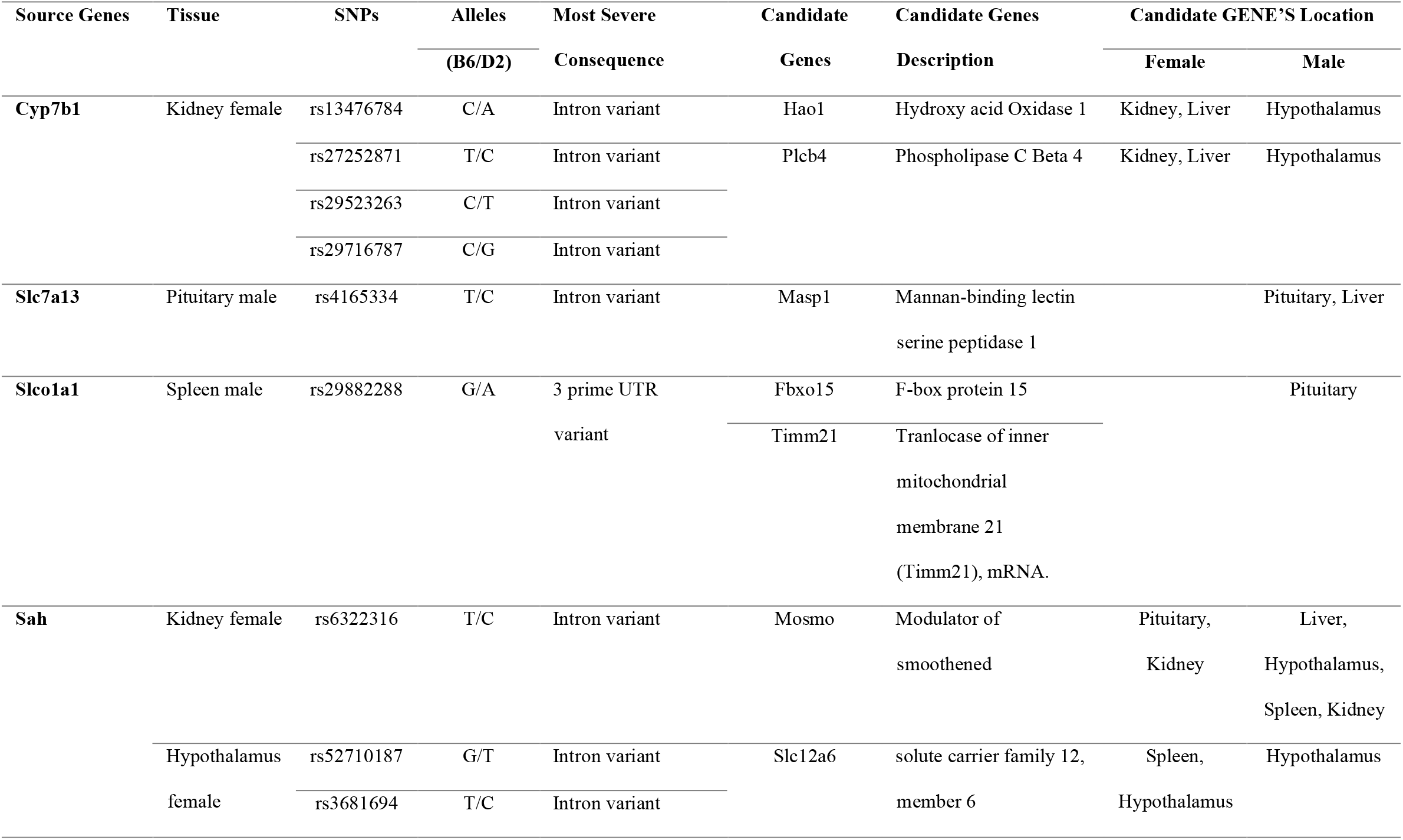
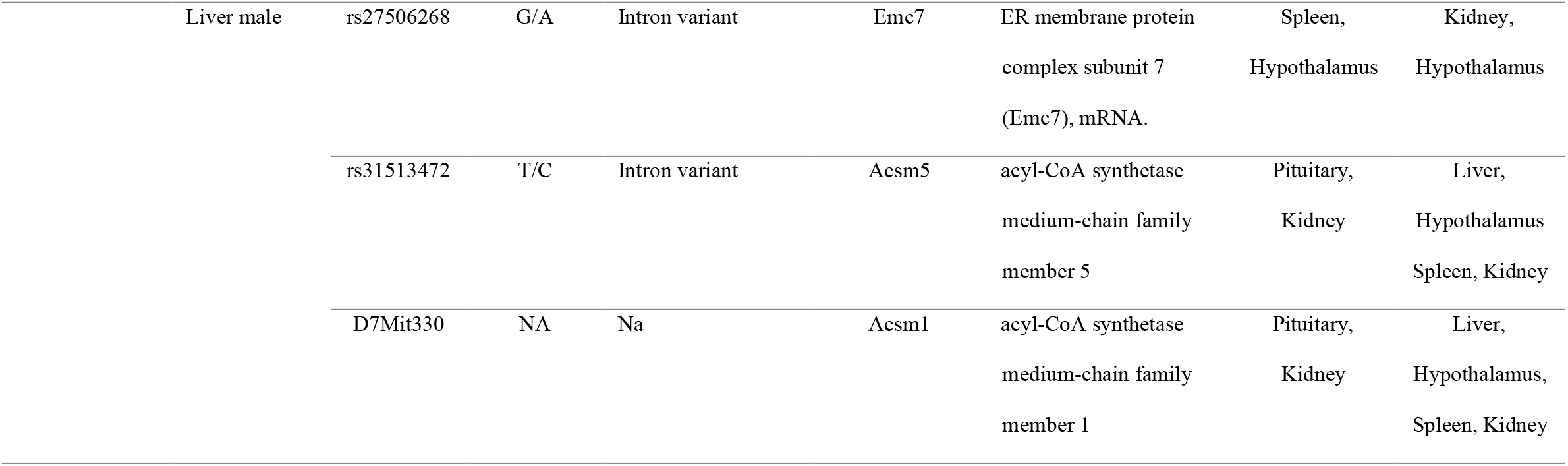
The information of SPNs and the candidate genes

The glycolate oxidase encoded by *Hao1* (Hydroxyacid oxidase 1, GCID: GC20M007863), is involved in the glyoxylate and dicarboxylate metabolic pathways, associated with human disorders primary hyperoxaluria and atopic dermatitis 3, respectively, and is also related to translation of viral mRNAs. *Hao1* is mostly expressed in the liver and pancreas. The phospholipase encoded by *Plc* (phospholipase C beta 4, GCID: GC20P009024) catalyzes the formation of inositol 1,4,5trisphosphate (IP3) and diacylglycerol (DAG) from phosphatidylinositol 4,5-bisphosphate. *Plc 4* is involved in the selective serotonin reuptake inhibitor and IP3 signaling pathways and DAG pharmacodynamics. (DAG), and IP3 signaling and is associated with human diseases of auriculocondylar syndrome 2 and uveal melanoma, respectively.

The candidate regulatory gene for *Scl7a13* in the pituitary gland and liver of male mice is *Masp1* (GCID: GC03M187216; Figure 3A), which encodes a serine protease that plays vital roles in the lectin complement pathway. The autosomal recessive disorder 3MC1 syndrome is associated with five mutations in *Masp1* (Atik et al., 2015). Examination of the correlations between *Scl7a13* and *Masp1* and their top related genes and phenotypes (Figure 3B) revealed that the gene most related to *Masp1*, Formin 1 (*Fmn1*), had negative correlations to the most related phenotype of *Slc7a13*, i.e., plasma lipidomics.

In the spleens of male mice, the significant e-QTLs of *Slcola1* included *Fbxo15* and *Timm21*. While the spleens of male B6 and D2 mice exhibited no statistically significant differences in expression of *Slco1a1* (*p* = 0.1672), its expression differs significantly in the pituitary gland of male mice (*p* = 0.0492) (Figure 3A). Correlation analysis suggests no strong correlation among *Slco1a1*, *Timm21*, and *Fbxo15* in the pituitary gland of male mice (Figure 3B).

The significant e-QTLs of *Acsm3* include *Acsm1* and *Acsm5* in the livers of male mice and *Mosmo* in the kidneys of female mice. *Mosmo*, *Acsm1*, and *Acsm5* localize to the same chromosome as Acsm3 and may serve as candidate cis-regulatory genes for *Acsm3* in the liver, hypothalamus, spleen, and kidney of male mice and the pituitary gland and kidney of female mice (Figure 3A).

The gene network between *Acsm3* and *Mosmo* showed that the top positively related phenotype of *Acsm3*, air control response, had strong negative correlations with *Mosmo* and its top positively related gene was serine/arginine-rich splicing factor (SRSF) kinase 2 (*Srpk2*; Figure 3B). *Acsm3*, also known as the *Homo sapiens* serine arginine (SA) hypertension-associated homolog (SAH), exhibited strong correlations with *Acsm1* and *Acsm5* (Figure 3B). *Acsm1* and *Acsm3* shared the same connection to negatively related phenotype *Streptococcus pyogenes* and their top negatively related genes exhibited strong positive correlations to each other. Crystallin mu (*Crym*), also known as NADP-regulated thyroid hormone binding protein (*Thbp*), was the top negatively related gene of *Acsm1*, which exhibited a medium negative correlation with *Acsm3* and a medium positive correlation with *Acsm5*.

The significant e-QTL of *Acsm3* in the hypothalamus of female mice included *Slc12a6* and *Emc7*, which may serve as candidate regulatory genes for *Acsm3* in the hypothalamus of male mice and the spleen of female mice. *Emc7* was also predicted to serve as a candidate gene for *Acsm3* in the kidney of male mice (Figure 3**.A9**). The correlations among *Acsm3*, *Slc12a6*, and *Emc7* showed that the top positively related phenotype of *Acsm3*, the MPTP (1-methyl-4-phenyl-1,2,3,6tetrahydropyridine) response, had medium negative correlations to *Emc7*. Similarly, the top positively related phenotype of *Acsm3*, GST (generalized seizure threshold) latency; Affy_10447921) was also the top negatively related gene of *Emc7*. In turn, *Emc7* had medium negative correlations to the top negatively related phenotype, chow diet metabolism.

#### 2.1.3 Rxfp1 differentially expressed and associated with genome expression in between sexes in Pituitary gland

The only DEG identified in the pituitary gland was *Rxfp1* (*p* = 0), which exhibited sex bias in pituitary gland, kidney and adrenal gland (Figure 4A). The top genes correlated to *Rxfp1* was *Akr1c14* (aldo-keto reductase family 1, member C14) in male mice and *Reln* in female mice. The top phenotype for *Rxfp1* in male mice was prodynorphin and in female mice was bone trait 38 (Figure 4B). *Rxfp1* in the adrenal gland and liver of female and male mice, respectively, exhibited a negative correlation to prodynorphin (*Pdyn*; Figure 4C). *Rxfp1* in the adrenal gland of female mice exhibited a moderate negative correlation to that in the hypothalamus of male mice, *Rxfp1* in the liver of male mice exhibited moderate negative correlation to that in the male kidney, and *Rxfp1* in the male hypothalamus exhibited moderate positive correlation to that in the female amygdala. *Rxfp1* (GCID: GC04P158315) encodes a member of the leucine-rich repeat-containing subgroup of the G protein-coupled 7-transmembrane receptor superfamily, which plays a critical role in sperm motility, pregnancy, and parturition.

**Figure 4.**
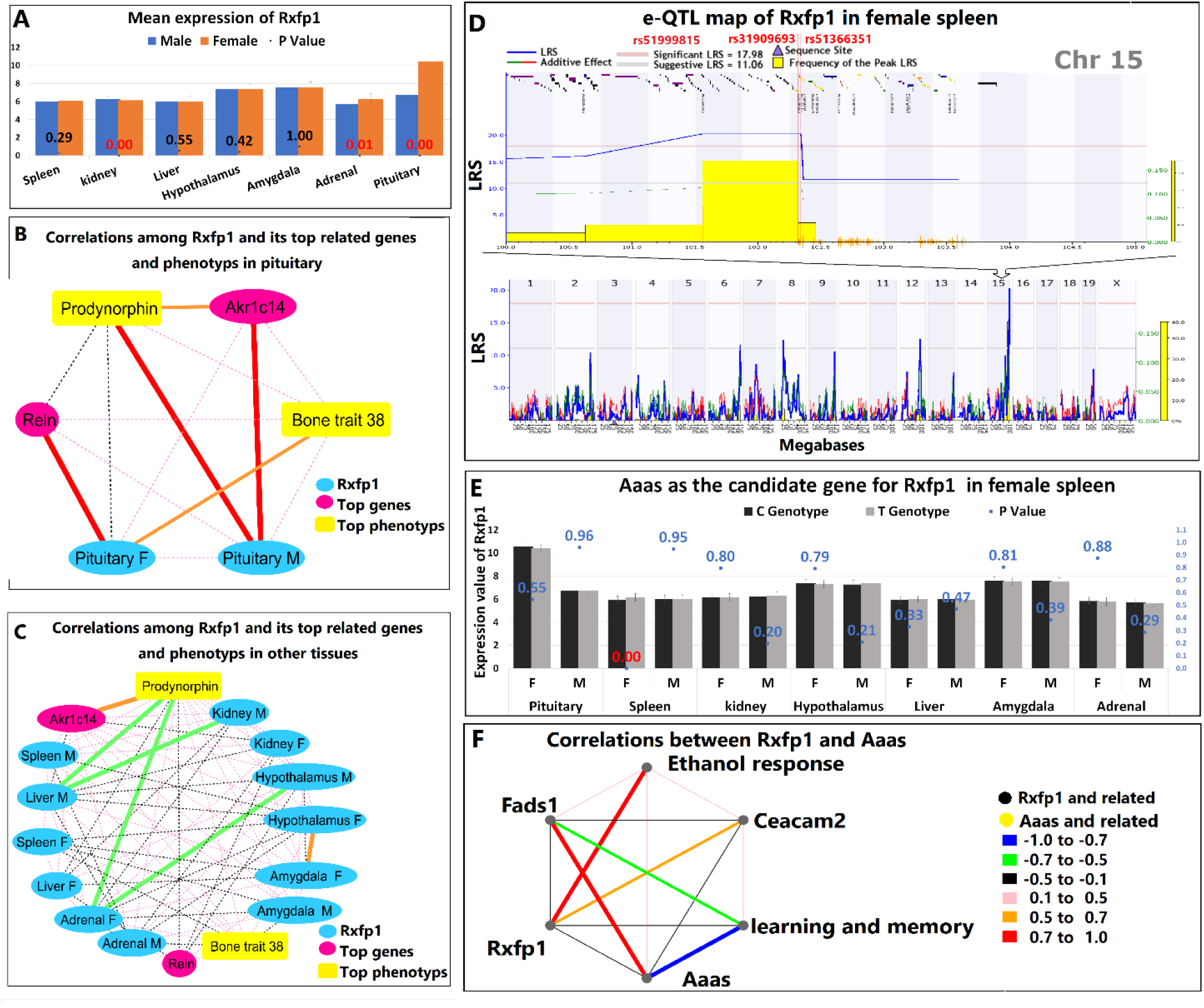
Expression of *Rxfp1*. **A.** The mean expression value of *Rxfp1* in all tissues examined. The blue and tangerine clustered column represents the mean expression of *Rxfp1* in male (blue) and female (tangerine) mice. The yaxis represents the expression values of *Rxfp1* in each tissue. The scatter markers represent the *p* values from the Student’s t-test; statistically significant differences (*p* < 0.05) are shown in red. There was sex differential expression of *Rxfp1* in pituitary gland, kidney, and adrenal gland. **B.** The networks of *Rxfp1* in the pituitary gland. The blue circles represent *Rxfp1*, the yellow rectangle represents the most related phenotypes to *Rxfp1* (Bone trait 38 in female mice and prodynorphin in male mice). The magenta circles represent the most related genes to *Rxfp1* (*Reln* in female mice and *Akr1c14* in male mice). We observed no strong correlations between expression of *Rxfp1* in male and female mice. **C.** The networks of *Rxfp1* in other tissues and its most related genes and phenotypes in pituitary gland. The blue circles represent *Rxfp1*, the yellow rectangles represent the most related phenotypes to *Rxfp1* in the pituitary gland (Bone trait 38 in female mice and prodynorphin in male mice), the red circle represents the most related genes to *Rxfp1* in the pituitary gland (*Reln* in female mice and *Akr1c14* in male mice). Prodynorphin had negative correlations to expression in the liver and adrenal glands of male and female mice, respectively. **D.** The bottom panel shows the e-QTL maps of *Rxfp1* in female spleen, while the top panel shows the e-QTL maps of *Rxfp1* on mouse Chr 15. The figure is labeled as described in the legend to Figure 1G, except that the vertical lines indicate the location of the SNPs, the SNPs that localize to the max LRS location and those that occur within genes are labeled in red, and the purple triangles on the x-axis indicate the location of *Rxfp1*. Three SNPs (red) mapped to the max LRS location and the frequency of peak LRS location and there were e-QTLs on mouse Chr 2 that were higher than the significant LRS level. **E.** The expression of *Rxfp1* between the B6 and D2 genotypes of *Aaas*. The panel is labeled as described in the legends to Figures 1A**, and** 3A. *Aaas* was identified as a candidate gene for Rxfp1 in the spleen of female mice. **F.** The correlations between *Rxfp1*, *Aaas*, and their most related genes and phenotypes in spleen. The black circle represents *Rxfp1* and its related genes and phenotypes, while the yellow circle represents *Aaas* and its related genes and phenotypes. We detected no strong correlation between *Rxfp1* and *Aaas*.

There were no statistically significant e-QTLs of *Rxfp1* in the pituitary gland, although one such *Rxfp1* e-QTL was mapped to Chr 15 in the spleen of female mice (Figure 4D). We identified three SNPs) and three candidate regulatory genes for *Rxfp1,* namely keratin 90 (*Krt90;* MGI:3045312), extra spindle pole bodies like 1 (*Espl1*) and aladin WD repeat nucleoporin (*Aaas*), as shown in **Supplementary table 19-20**. Krt90 has no human homolog and will not be considered further here.

Expression of *Espl1* was controlled by a single SNP, rs31909693, whose most abundant server sequence is a non-coding C/A transcript exon variant. The genome of the B6 mouse strain has a C nucleotide at this position, while that of the D2 strain has an A at this position. Similarly, expression of *Aaas* is controlled by a single SNP, rs51366351, whose most abundant server sequence was intron variant C/T. The genome of the B6 mouse strain has a C nucleotide at this position, while that of the D2 strain has a T at this position. According to the approach described by Tabor *et al*. (2012), the candidate regulatory gene for *Rxfp1* may be *Aaas* (Tabor et al., 2002). The expression value of the C and T genotypes mapped for *Rxfp1* suggested that *Aaas* is a candidate regulatory gene for *Rxfp1* in the spleen of female mice (Figure 4E). There were no strong correlations among *Rxfp1*, *Aaas*, or their top related genes and phenotypes (Figure 4F).

#### 2.1.4 The sex bias genes in spleen are crusted to heme biosynthesis network and may contribute to the difference hemoglobin level between male and females

We identified 26 (0.09%) sexually dimorphic genes in spleen (FDR<0.05) but none of them were predicted to have *p* = 0 (Table 1). STRING network analysis of these genes suggested the existence of a network that was clustered as heme biosynthesis and iron import into the mitochondrion (**Supplementary figure E).** Three genes, the probably mitochondrial glutathione transporter (*Slc25a39*), solute carrier family 25 member 37 (*Slc25a37*), and ferrochelatase (*Fech*), which play important roles in biosynthesis of heme, were clustered as seed genes. Slc25a39 and Slc25a37 are solute carrier family 25 members, while Fech encodes ferrochelatase. The mean expression values revealed that all three genes were expressed at higher levels in male mice than in female mice (**Supplementary figure F).**

#### 2.1.5 Liver has the most sexually differential expressed genes other than sex organs and these genes contribute to the metabolism functions between males and females

Liver is the major site for hormone metabolism that is sexually dimorphic (Langer and Chiandussi, 1987). Our analysis revealed 121 genes in mouse liver that were differentially expressed between males and females (**Supplementary table 21**). Analysis of the mean expression levels of these DEGs revealed that 87 of them were expressed at higher levels in male mice than in female mice (Figure 5A). Many of these differences are known to be driven by the pituitary gland-derived growth hormone (GH), which acts through signal transducer and activator of transcription (STAT) 5b and hepatocyte-enriched nuclear factor (HNF) 4α. STAT-5b and HNF-4α directly and indirectly regulate expression of many steroid and xenobiotic metabolizing enzymes, leading to sexually dimorphic expression in several important biological pathways (Gatti et al., 2010).

**Figure 5.**
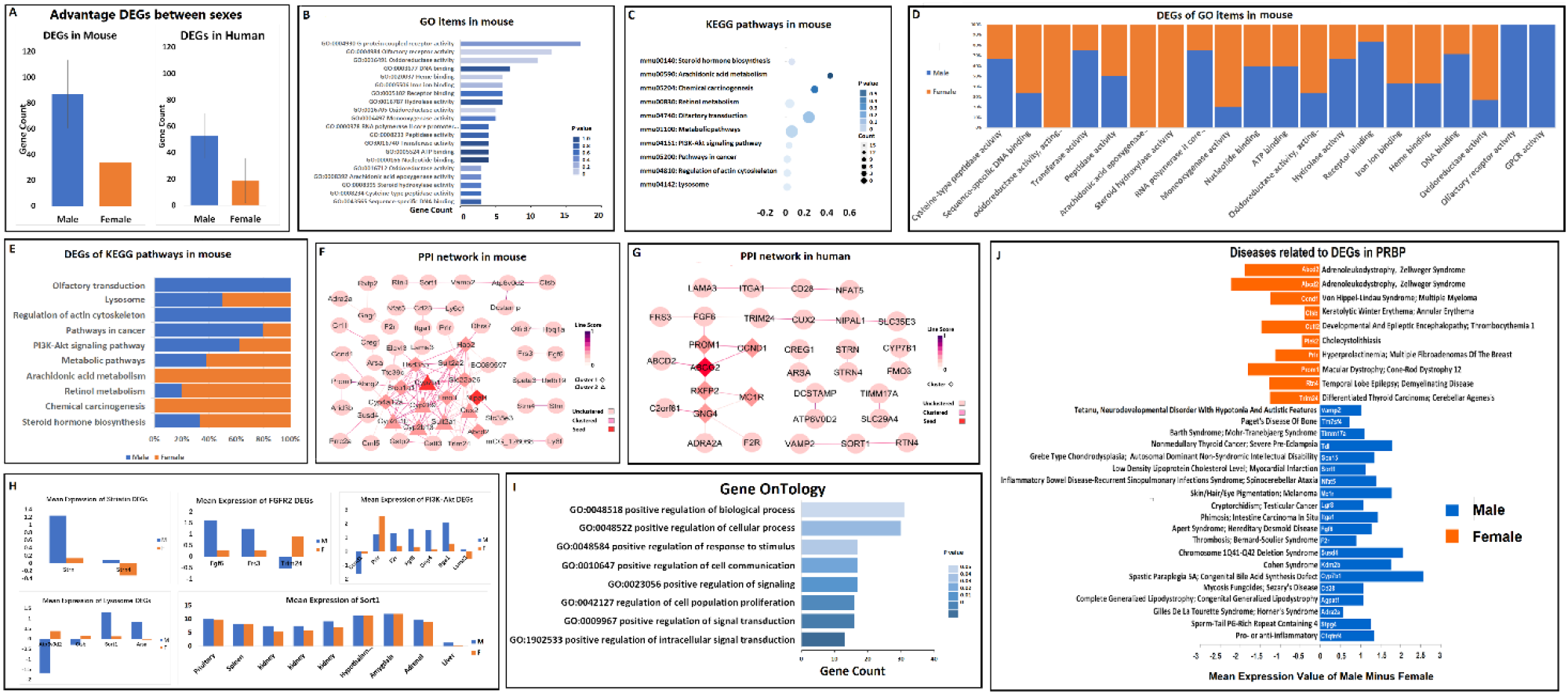
Comparison of DEGs in mouse and human tissues. **A.** The advantage DEGs in mouse and human tissues. The y-axis represent the number of DEGs. In both mouse and human tissues, more sex DEGs were expressed at higher levels in males than in females. **B.** The top 20 GO terms clustered by function annotation. The x-axis represent gene numbers of each GO term, while the y-axis represent the GO terms in the order of low to high. The colors of the GO terms from light blue to dark blue represent their *p* values from low to high. **C.** The volcano plot of KEGG pathways of DEGs in mouse. The x-axis represent the *p* value, the y-axis represent the different KEGG pathways, and the size of the circle reflects the gene counts. **D.** The column of advantage DEGs in different GO terms. The x-axis represent the GO terms and the y-axis represent the proportions of advantage genes expressed in males and females. Males exhibited advantages in GPCR activity and olfactory receptor, while females exhibited advantages in arachidonic acid epoxygenase activity and steroid hydroxylase activity. **E.** The column of advantage DEGs in KEGG pathways. The x-axis represents the proportions of advantage gene expressed in males and females, while the y-axis represent KEGG pathways. Males exhibited an advantage in olfactory receptor activity pathways and females exhibited advantages in chemical carcinogenesis and arachidonic acid metabolism. **F.** The protein-protein interaction (PPI) network of DEGs in mice. The pink nodes represent the unclustered DEGs, the rosy-red nodes represent the clustered DEGs, and the red nodes represent seed DEGs. The colors of the lines from light to dark represent the correlation values from zero to one. The triangle shaped nodes represent cluster one and the diamond shaped nodes represent cluster two. *Cyp7b1* and *Nipal1* were clustered as seed genes. **G.** The PPI network of DEGs in humans. The panel is labeled as described for Figure 5F, except that the diamond shaped nodes represent clustered genes. *Abcg2* was clustered as a seed gene. **H.** Mean expression values between sexes in different pathways or clusters. The x-axis represents the gene name and the y-axis represents the mean expression values of genes that were modified by at least log2-fold. The blue columns represent males, while the orange column represents females. **I.** The human GO terms clustered by function annotation. The x-axis represents the numbers of genes for each GO terms and the y-axis represents the GO terms in the order of low to high. The colors of the GO terms from light blue to dark blue represent the *p* values from low to high. **J.** The DEGs related to biological process. The genes in the orange columns represent higher expression in females than in males, while the genes in the blue column represent higher expression in males than in females. The x-axis represents the mean expression value of the indicated genes; genes expressed at higher levels in males are shown as negative values, while genes expressed at higher levels in females are shown as positive values. The y-axis shows the name and description of each gene.

We used functional annotation clustering to study these liver DEGs further. The top 20 GO terms (Figure 5B) and the top10 KEGG pathways (Figure 5C) were clustered and listed as a function of their p values. Each DEG was examined for its proportion of advantage in male and female mice (Figure 5D-5E), The results suggest that male mice had the advantage in GPCR activity, olfactory receptor, and the olfactory receptor activity pathways, while female mice had the advantage in arachidonic acid epoxygenase activity, steroid hydroxylase activity, chemical carcinogenesis, and arachidonic acid metabolism.

To better understanding the interactions between and among these DEGs, we used the STRING database to construct a protein-protein interaction network (PPI) and visualized the network using Cytoscape software (Shannon et al., 2003), as shown in Figure 5F. We found that the female advantage DEGs were predicted to predominantly play roles in metabolism and oxidation-reduction reactions. These DEGs included the mouse genes flavin-containing dimethylaniline monooxygenase 3 (*Fmo3*), hydroxy acid oxidase 2 (*Hao3*), sulfotransferase family 2A member 1 (*Sultx2*), sulfotransferase family 2A, dehydroepiandrosterone (DHEA)-preferring, member 2 (*Sult2a2*), carbonyl reductase [NADPH] 1 (*Cbr1*), glutathione S-transferase theta-3 (*Gstt3*), and cytochrome P450, family 2, subfamily b, polypeptides 9 (Cyp2b9), 10 (Cyp2b10), and 13 (Cyp2b13). Metabolic enzymes contribute to detoxification of biotransformation intermediates and Phase II enzymes participate in conjugation and inactivation of chemical carcinogens in oxidation reactions. Most of these enzymes are expressed at higher levels in female mice, which might confer on them the advantage of a lower incidence of carcinomas (Islami et al., 2021; Oliveira et al., 2007). Interestingly, the expression of many of these genes were altered in growth hormone-releasing hormone (GHRH) knockout (KO) mice, which exhibit a significantly increased maximum lifespan in both males and females due to the GHRH deletion (Sun et al., 2013). It is unknown whether differential expression of these genes also contributes to the longer life expectancy in females.

In male mice, DEGs related to proteinase activity and the regulation of the signaling pathways were expressed at higher levels, including, fibroblast growth factor 6 (*Fgf6*), integrin subunit alpha 1 (*Itga1*), laminin subunit alpha 3 (*Lama3*), MINDY lysine 48 deubiquitinase (*Mindy3*), and caspase 14 (*Casp14*). Expression of G-protein coupled receptor genes including MAS-related GPR family member E (*Mrgpre*), melanocortin 1 receptor (*Mc1r*), relaxin/insulin-like family peptide receptor 2 (*Lgr8*), coagulation factor II thrombin receptor (*F2r*), and odorant receptor genes (*Olfr*s) was also higher in male mice, perhaps due to sex-specific cell signaling (Valentino et al., 2013). The cytoskeleton and cell movement are required in physiological processes of inflammation, immune response, wound healing, and cancer development. Sex steroids can activate sex steroid receptors to recruit intracellular molecular cascades that include phosphatidylinositol 3 kinase (PI3K) to regulate the actin cytoskeleton (Giretti and Simoncini, 2008). Therefore, these DEGs may result from kinase cascades signaling subsequent to binding of sex steroids to their cognate receptors.

We used the STRING database to explore the differences in expression of these genes between mice and humans. Seventy of these DEGs are also expressed in humans (Figure 5A**, Supplementary table 22**). PPI network analysis (Figure 5G) allowed identification of one protein domain, one local network cluster, two KEGG pathways, and eight biological processes (**Supplementary table 23**). These human-relevant proteins are described in detail below.

##### Protein domain: Striatin family

Two DEGs that exhibited higher expression values in male mice than female mice (Figure 5H) were identified as members of the striatin family. Striatins are scaffolding proteins that possess multiple protein-binding domains to facilitate their interaction with and activation of signal transduction molecules. For example, the calcium–calmodulin (CaM)binding domain is involved in glutamate regulation pathways and may play a crucial role in the cellular mechanisms of learning and memory (Benoist et al., 2006). Ruhs *et al*. (2017) showed that striatins play a role in signaling through the plasma membrane mineralocorticoid receptor (MR) and estrogen receptor (ER) (Ruhs et al., 2017). Striatin expression is regulated by a feedback loop in which its direct binding to estrogen receptor alpha (Erα) in female mice reduces levels of striatin protein. In turn, this may serve to protect the mice from the salt-sensitive elevation of blood pressure (Garza et al., 2015; Gupta et al., 2017).

##### Local network cluster: Activated point mutants of fibroblast growth factor-activated receptor 2 (Fgfr2) and the effect on FGFR2 activity

Fibroblast growth factor 6 (*Fgf6*; GCID:

GC12M004413) and other FGF family members are involved in biological processes that include embryonic development, cell growth, morphogenesis, tissue repair, and tumor growth and invasion. Activated *Fgfr1* point mutations are involved in a variety of cancers (Wu et al., 2013). In the current study, we identified three DEGs whose mean expression vary between male and female mice (Figure 5H). Fibroblast growth factor receptor substrate 3 (*Frs3*; GCID: GC06M043574) links FGF to downstream signaling pathways that involve activation of mitogen-activated protein kinases (MAPKs). Specific expression of tripartite motif-containing 24 (*Trim24*), also known as transcription intermediary factor one alpha (*Tif1α*), may contribute to the decreased incidence of hepatocellular carcinoma, although the balance of its functions as tumor suppressor versus oncogene is species- and tissue-specific (Appikonda et al., 2016). Higher expression values of *Fgf6* and *Frs3* and a lower expression value of *Trim24* in male mice may cause them to develop cancer more readily than do female mice.

##### KEGG Pathway: PI3K-Akt signaling

The *Pi3k-Akt* (protein kinase B) pathway is regulated by growth factors and its diverse downstream effects on cellular metabolism occur through either direct regulation of nutrient transporters and metabolic enzymes or via the control of transcription factors that regulate the expression of key metabolic pathway components. The Pi3k-Akt pathway is the most commonly activated pathway in human cancers (Hoxhaj and Manning, 2020). In the current study, we identified seven DEGs that were predicted to be members of this pathway, only two of which were expressed at higher levels in female mice (Figure 5H), specifically the prolactin receptor (*Prlr*) that is responsible for transducing prolactin signaling (Pujianto et al., 2010) and cyclin D1 (*Ccnd1*), an important cell cycle-dependent regulator of cellular proliferation. The central role of *Ccnd1* in the pathogenesis of cancer is illustrated by its driving of unchecked cellular proliferation that promotes tumor growth when it is expressed at high levels (Montalto and De Amicis, 2020). The sex-specific difference in *Ccnd1* expression was identified in rat liver (Kwekel et al., 2017) but it remains unknown whether *Ccnd1* contributes to sex differences in the genesis and progression of some human cancers. Pi3k-Akt can also activate fusion of the mTOR complex 1 (mTORC1) protein with lysosomes during autophagy, which is involved physiological and pathological processes that are closely related to tumorigenesis (Yu et al., 2018).

##### KEGG Pathways: the lysosomal pathway

We identified four DEGs that are enriched for lysosomal pathway functions (Figure 5H). Sortilin (*Sort1*), also known as neurotensin receptor 3 (*Ntr3*), serves as a multi-ligand sorting receptor that plays important roles in endocytosis and intracellular protein trafficking (Musunuru et al., 2010). In male mice, *Sort 1* is expressed at higher levels in both liver and kidneys. Although the mechanisms that regulate *Sort1* expression and function remain unclear, deregulation of *Sort1* function has been implicated in four major human diseases: cardiovascular disease, type 2 diabetes, Alzheimer’s disease, and cancer (Wilson et al., 2016). More research will be needed to determine whether the sexually dimorphic expression of *Sort1* contributes to the development of diseases that differ in incidence and progression between males and females. The protein encoded by the *Arsa* (arylsulfatase A; GCID: GC22M050622) gene hydrolyzes cerebroside sulfate to cerebroside and sulfate. Defects in this gene lead to the autosomal recessive disorder metachromatic leukodystrophy (MLD), although there have been no descriptions of sex bias associated with this disease. Also identified were damage-associated microglia (DAM) genes *Ctsb* (cathepsin B) and *Atp6v0d2* (ATPase H^+^ transporting V0 Subunit D2) that play roles in the development of Alzheimer’s disease (Srinivasan et al., 2019).

##### Biological Processes (GO terms)

Eight GO terms are associated with positive regulation of biological processes (PRBP) (Figure 5I). Gene regulatory processes serve as drivers of sex-specific differences in human health and diseases (Lopes-Ramos *et al*., 2020). The current study identified 30 DEGs (10 in female mice and 20 in male mice) in this category, whose related diseases and mean expression value are shown in Figure 5J. Among them, *Ccnd1*, *Cyb7b1*, *Trim24*, and reticulon-4 (*Rtn4*) are controlled by GH (Sun *et al*., 2013). Four other genes, *Prlr*, *Lgr8*, *Itga1*, and sperm-tail PG-rich repeat containing 4 (*Stpg4*) are related to diseases of the reproductive organs in male or female mice. The rest of this group of DEGs are associated with diseases characterized by wellrecognized sex-biased manifestations, including fat biogenesis, autoimmune thyroid disease, neurological disorders, and cardiomyopathy (Lopes-Ramos *et al*., 2020).

### 2.2 Sex differences on the X chromosome

In detecting differential expression (q < 0.05) of the X chromosome, the proportion of probes that allowed analysis in the tissues of interest and the numbers of identifiable unique genes were as follows (Table 1): adrenal gland, 23.53% (284 of 1,207 probes); amygdala, 0.83% (10 of 1,207 probes); hypothalamus, 0.83% (10 of 1,207 probes); kidney, 0.21% (3 of 1,442 probes); liver, 28.65% (198 of 691 probes); pituitary gland, 20.63% (249 of 1,207 probes); and spleen, 1.02% (13 of 1,271 probes). This distribution indicates that X chromosome genes that exhibit differential expression were predominantly found in the adrenal gland, liver, and pituitary gland, while the number of DEG on the X Chr were low in hypothalamus and kidney.

Some genes exhibited statistically significant differences in expression intensities between the sexes. We were particularly interested in the genes whose differential expression exhibited very low p values (*p* = 0), namely DEAD-box helicase 3 X-linked (*Ddx3x*) in amygdala, eukaryotic initiation factor 2 subunit 3, X-linked (*Eif2s3x*), lysine demethylase 5C (*Kdm5c*), histone lysine demethylase 6A (*Kdm6a*) and *Xist*. The tissue specificity of these DEGs was as follows: amygdala, *Ddx3x*, *Eif2s3x*, *Kdm5c*, and *Xist*; hypothalamus, *Ddx3x*, *Eif2s3x*, *Kdm5c*, *Kdm6a*, and *Xist*; kidney, *Xist*; spleen, *Eif2s3x*, *Kdm6a*, and *Xist*; adrenal gland, 11 genes including *Eif2s3x*, *Kdm5c*, *Kdm6a*, and *Xist*; liver, 13 genes including *Eif2s3x*, *Kdm6a*, and *Xist*; and pituitary gland, 17 genes including *Ddx3x*, *Eif2s3x*, *Kdm5c*, *Kdm6a*, and *Xist* (**Supplementary table 24**). The mean expression of the DEGs on the X Chr is shown in Figure 6A. Strikingly, the *Xist* gene exhibited higher expression in all tissues examined in female mice relative to that of male mice.

**Figure 6.**
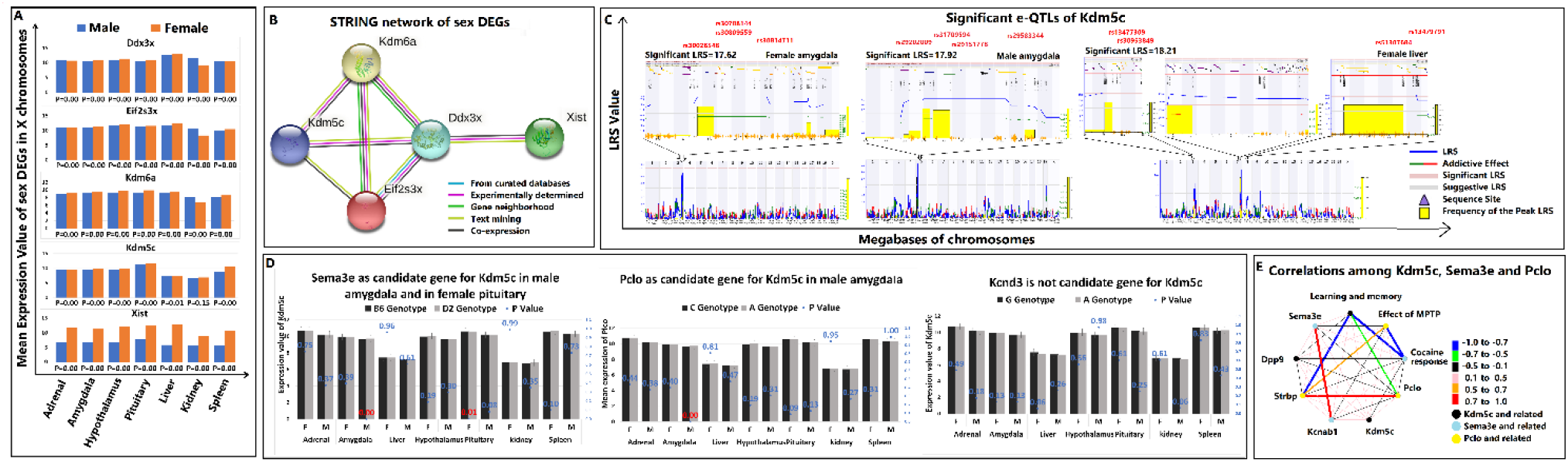
Sexually dimorphic expression of genes on the X chromosome. **A.** The mean expression value of X Chr DEGs in all tissue. Except for *Kdm5c* in kidney, all genes exhibited sex bias in all tissues we examined (P < 0.05 for each). **B.** The STRING networks of X Chr DEGs. The colored nodes represent query proteins and the first shell of interactors, while the edges represent protein-protein associations **C.** The e-QTL maps of *Kdm5c*. The bottom panel shows the interval map of the entire genome sequence and the upper panel shows the map within the significant QTL location. The panel is labeled as described in the legends to Figures 1G and 4D, except that the purple triangles represent the location of each DEG. The image on the left shows the e-QTL map of *Kdm5c* in female amygdala, for which none of the SNPs localized to the peak LRS location. The image in the middle shows the e-QTL map of *Kdm5c* in male amygdala, for which all SNPs localized to the peak LRS location. The image on the right shows the e-QTL maps of *Kdm5c* in female liver, in which three mouse chromosomes had significant e-QTLs: mouse Chr 3, Chr 7, and Chr 8. Only SNP rs13477309 on Chr 3 localized to the peak LRS location. **D.** Differences in *Kdm5c* expression of the candidate genes between B6 and D2 mouse genotypes. The panel is labeled as described in the legends to Figures 1A and 3A. The title of each panel indicates the tissue localization of the candidate genes. **E.** The correlations among *Kdm5c* and its candidate regulator genes. The black circles represent *Kdm5c* and its top related genes and phenotypes, the blue circles represent *Sema3e* and its top related genes and phenotypes, the yellow circles represent *Pclo* and its top related genes and phenotypes, and the colored edge represents the sample correlation.

The interactions among the Chr X DEGs examined by STRING network (Figure 6B) revealed complex interaction patterns. However, the functional enrichment highlighted a common molecular function, namely nucleic acid binding. Despite the DEGs sharing localization to the X Chr and similar functions, their protein products share no more 10% amino acid identity and no more than 23% amino acid similarity (**Supplementary figure D**), which is well below the 40% sequence identity required for predicted similar functionality (Pearson, 2013). Thus, the proteins encoded by the X Chr DEGs are likely to present different functions and participate in different aspects of common biological processes.

We also identified DEGs in both autosomal and X chromosomes that exhibited no meaningful similarities in protein products prompted us examine proteins encoded by genes that do not exhibit sexually dimorphic expression (non-DEGs). Proteins encoded by non-DEGs on the X Chr exhibit less than 11% amino acid identity and less than 24% amino acid similarity, consistent with the levels of amino acid identity and similarity among proteins encoded by DEGs on autosomal and X chromosomes (**Supplementary figure D**). This finding argues that the emergence of DEGs in autosomal and X chromosomes is unrelated to amino acid commonalities among encoded proteins or that the genetic regulatory elements that respond to sex-specific regulators are likely to be located outside protein coding regions.

The X chromosome-linked DEGs mapped to significant e-QTLs only for *Kdm5c* in the liver and hypothalamus of female mice and the hypothalamus of male mice (Figure 6C). Combining the interval analysis with the e-QTL map (Figure 6C**, Supplementary table 25-27**), we identified three candidate regulatory genes for *Kdm5c*, namely semaphorin 3E (*Sema3e*) on SNPs rs29202009 and rs31709594, piccolo presynaptic cytomatrix protein (*Pclo*) on SNP rs29151778, and potassium voltage-gated channel subfamily D member 3 (*Kcnd3*) on SNP rs13477309. Comparison of the *Kdm5c* expression values of candidate SNPs between the B6 and D2 mouse genotypes (Figure 6D), identified *Sema3e* as a candidate regulatory gene for *Kdm5c* in pituitary gland of female mice and both *Sema3e* and *Pclo* as predicted *Kdm5c* candidate regulatory genes in the amygdala of male mice. The correlations among *Kdm5c*, *Sema3e*, and *Pclo* (Figure 6E) show that the top related phenotype of *Kdm5c* (learning and memory; L&M) had a strong negative correlation to *Pclo* and its top related gene. The top related phenotype of *Sema3e*, cocaine response, had a negative correlation with L&M and with the top related phenotypes of *Pclo*, effect of 1-methyl-4-phenyl-1,2,3,6-tetrahydropyridine (MPTP), which causes symptoms of Parkinson’s disease.

## 3. Discussion

### 3.1 Genes on autosomal chromosomes that show sexually dimorphic expression

***Akr1d1*** is essential for bile acid biosynthesis and inactivation and clearance of steroid hormones (Barnard et al., 2020). *Akr1d1* expression in the liver of female was higher than that in male mice (*p* = 3.28E-06), as shown in Figure 1B. *Akr1d1* knockdown in cultured human cells increased hepatocyte accumulation of triglycerides, sensitivity to insulin, and glycogen synthesis through increased *de novo* lipogenesis (DNL) and decreased β-oxidation, thereby fueling inflammation in hepatocytes (Nikolaou et al., 2019). This may be related to the higher incidence of non-alcoholic fatty liver disease (NAFLD) in men relative to that in women (Rich et al., 2018). *Adr1d1* KO and *Adr1d1*-difficient female mice are more protected from the adverse metabolic effects of a high-fat diet than are male mice (Phelps et al., 2019). Moreover, the observed higher expression of *Adr1d1* in female non-human primates is meaningful in light of the estrogen-induced increase in canalicular secretion of cholesterol into the bile, which can help to eliminate excess steroid hormone (Colvin Jr et al., 1998).

***Cyp7b1*** protein is localized to the ER membrane, where its expression is positively regulated. by activation of ER-α. The *Cyp7b1* gene encodes a cytochrome P450 superfamily enzyme that participated in the alternative pathway of bile acid synthesis from cholesterol (Phelps *et al*., 2019) and played roles in the metabolism of endogenous oxysterols and steroid hormones (Stiles et al., 2009).The higher expression value of *Cyp7b1* in male mice (Figure 2B) suggests that the alternative pathway of bile acid synthesis may be more active in male mice than in female mice, perhaps resulting from a greater number of methylation sites in *Cyp7b1* proteins in male mice than in female mice (Penaloza et al., 2014). The diseases related to *Cyp7b1* mutations are mainly associated with cholesterol metabolism and bile acid synthesis, which can cause neonatal liver disease and motorneuron degenerative disease, although the existence of sex-related differences in these diseases remains unknown (Chen et al., 2020; Tsaousidou et al., 2008).

***Slc7a13*** (Solute Carrier Family 7 Member 13, GCID: GC08M086214) encodes a transmembrane protein that mediates the transport of L-aspartate and L-glutamate in a sodiumindependent manner (Matsuo et al., 2002) and associates with developmental and epileptic encephalopathy 3 and hyperekplexia 3 (Yahyaoui and Perez-Frias, 2019). *Slc7a13* is related to retinitis pigmentosa and higher *Slc7a13* expression in male mice and humans may contribute to a higher incidence of retinal degeneration (Jin et al., 2014; Li et al., 2019). ***Slco1a1*** (solute carrier organic anion transporter family, member 1a1), also known as *Oatp1* (ornithine aminotransferase pseudogene 1, GCID: GC10M124443), is a pseudogene in humans. It encodes a transmembrane protein that mediates the sodium-independent transport of organic anions, transporting endogenous substances including cyclic guanosine monophosphate (cGMP), bile acids, and hormone derivatives (Thakkar et al., 2017). One of the *Slco1a1*drug substrates is methotrexate (MTX), which exhibits differential pharmacological patterns in men and women. Higher expression of *Slco1a1* in men may contribute to a lower incidence of adverse events leading to MTX modifications than is observed in women (Giusti et al., 2019).

***Acsm3*** (acyl-CoA synthetase medium-chain family member 3; GCID: GC16P020610), also known as ***Sah*** (SA rat hypertension-associated homolog) and other members of the acyl-CoA synthetase medium chain family that also includes *Acsm1* and A*csm5* are related to fatty acid betaoxidation pathways in peroxisomes and mitochondria and catalyze the production of acyl-CoA, the first step in fatty acid metabolism, from fatty acids and coenzyme A (Iwai et al., 2002; Vessey et al., 1999). *Acsm3* and candidate *cis*-regulatory genes *Mosmo*, *Acsm5*, and *Acsm1* are all located on mouse Chr 7. *Mosmo* (Modulator of Smoothened, GC16P022007) plays a role in the hedgehog signaling pathway that transmits information essential for proper cell differentiation to vertebrate embryonic cells. The aberrant reactivation of hedgehog signaling pathway after development is associated with malignant skin, brain, prostate, breast, and hematological (blood) cancers (Montagnani and Stecca, 2019). The higher expression of *Acsm3* we observed in male mice may reflect greater activation of the hedgehog signaling pathway related to higher incidences of cancer in male vertebrates, including humans.

***Rxfp1*** encodes a G-protein coupled receptor that is regulated by the hormone relaxin, leading to stimulation of adenylate cyclase and a concomitant increase in cAMP levels. As such, *Rxfp1* plays a critical role in sperm motility, pregnancy, and parturition. *Rxfp1* was expressed at higher levels in mouse kidney but at lower levels in adrenal and pituitary glands. Higher expression of *Rxfp1* in the kidney of male mice may suggest a connection between sperm motility and kidney function in men (Lundy and Vij, 2019).

### 3.2 Genes on sex chromosomes that show sexually dimorphic expression

***Xist*** (X-inactive specific transcript) is a regulatory RNA gene that affects the variation in X chromosome gene expression between sexes and has been used for chromosome therapy (Balaton et al., 2018; Crowley *et al*., 2015). The expression of *Xist* at higher levels in female mice than in male mice (Figure 6A) is reasonable since the process of X-Chr-inactivation occurs in the female X chromosome inactivation center. *Xist* is an oncogene that accelerates tumor progression and is related to the hypogonadotropic hypogonadism (HH) described in cryptorchid male children (Hadziselimovic et al., 2017; Li et al., 2018).

***Kdm5c* (**Lysine (K)-specific demethylase 5C) encodes an enzyme that demethylates lysine residues in the N-terminal tail of histone H3. The enzyme encoded by ***Kdm6a*** (Lysine(K)-specific demethylase 6A) also demethylates lysine residues in the N-terminal tail of histone H3 but its target lysine differs from the one targeted by *Kdm5c*. Both are X-linked chromatin-modifying genes whose protein products regulate transcription of eukaryotic genes (Tricarico et al., 2020). Mutation of *Kdm5c* causes X-linked inherited intellectual disability that is observed at a higher rate in males than in females, as most heterozygous female carriers are asymptomatic (Schenkel et al., 2018). *Kdm6a* variants cause X-linked Kabuki syndrome type 2 and are more likely have maternallyinherited variants (Faundes et al., 2021).

***Eif2s3x*** (eukaryotic translation initiation factor 2, subunit 3, structural gene X-linked) encodes a subunit of eukaryotic initiation factor 2 (eIF2), which is involved in the early steps of protein synthesis. Mutations in *Eif2s3x* are associated with the severe X-linked intellectual disability syndrome MEHMO (mental retardation, epileptic seizures, hypogonadism, microcephaly, obesity), of which all reported cases have been in boys (Skopkova et al., 2017).

***Ddx3x*** (DEAD-box helicase 3, X-Linked) encodes a multi-domain protein that plays key roles in the nucleus and the cytoplasm, including transcriptional regulation, mRNP assembly, pre-mRNA splicing, mRNA export, translation, cellular signaling, and viral genome replication (Cargill et al., 2021). The downregulation of *Ddx3x* expression in hepatocellular carcinoma (HCC) is more frequently observed in men than in women, perhaps in part because HCC is more common in men (Chang et al., 2006). The interindividual variability in *Ddx3x* expression potentially could be associated with differences in cancer predisposition among women (Garieri et al., 2018). Moreover, Mutations in *Ddx3x* have also been associated with unexplained intellectual disability (Snijders Blok et al., 2015).

All five of these genes have been associated with escape from epigenetic inactivation of one XChr, which would normally result in its silenced transcription, and contribute to sexually dimorphic traits and sex bias in cancer (Balaton *et al*., 2018; Dunford et al., 2017). Mutations in many X-linked genes have been associated with intellectual deficiency. Because XX female embryos receive two X chromosomes, one each from their mother and father, XX females are biologically more susceptible to X-linked disorders than are XY males (Migeon, 2020).

### 3.3 The differences and correlations of DEGs genes across tissue

#### 3.3.1 Different organs show variability in sex dimorphism

The seven tissues we examined relate to four body systems, namely the endocrine system (the adrenal gland and the anterior lobe of the pituitary gland), the lymphatic system (spleen), the digestive system (liver), the nervous system (amygdala, hypothalamus, and the posterior lobe of the pituitary gland), and the urinary system (kidney). The proportion of DEGs in different chromosomes were mapped in Figure 7A. To summarize our observations, **1)** the liver contained the most sex DEGs of the organs we examined, but the proportion of DEGs in the liver vary least from chromosome to chromosome. **2)** The kidney was the organ we examine that seemed least affected by sexual dimorphism of gene expression, as the proportion of DEGs in any chromosome was not very high and these proportions did not vary widely. This observation may provide at least a partial explanation for sex being an impact factor for female-to-male transplantation of liver but not kidneys (Morgan et al., 2020; Uno et al., 2017). **3)** Our examination of nervous system tissues were limited to the amygdala and hypothalamus, in which we observed no sexual dimorphism of expression for genes on autosomal chromosomes. This result may suggest that for any observed differences between males and females in the activity of brain neuronal circuitry within the amygdala and hypothalamus, the sex chromosomes may represent independent impact factors. For example, many X-linked genes are associated with intellectual deficiency (Migeon, 2020) and it has been suggested that the differences between males and females in cognitive processing such as language learning (Scheuringer *et al*., 2017) or in stress behaviors (Donner and Lowry, 2013) are more inherited and would be difficult to be modified directly by the environment. **4)** The pituitary gland exhibited similar patterns to those observed in the adrenal gland but had fewer sex differential genes (Table 1, Figure 7). The pituitary gland is regarded as an important representative of endocrine organs due to its dual origin from oral ectoderm (anterior lobe) and neural ectoderm (posterior lobe) and the secretion of the majority of hormones by its anterior lobe (Hong et al., 2016). **5)** We observed that the spleen contained more sex differential genes on sex chromosomes than did the kidney or brain but far fewer than did the endocrine organs or liver. The sex differential genes in spleen were mainly related to erythropoiesis (**Supplementary figure E**). This finding may help to explain why the number of red blood cells per microliter of blood is higher in men than in women. Elevated levels of androgenic hormones like testosterone in men may be related to increased blood hemoglobin levels, which may lead to increased physical strength in males (Handelsman et al., 2018). Interestingly, observed differences in the weight of the spleen, which tends to be greater in men, are eliminated after gonadectomy (Masui and Tamura, 1926). A recent study showed that sexual dysfunction in both men and women increased after splenectomy(Sabino and Petroianu, 2021). These findings suggest that sex hormones may play an important role in the anatomy and function of the spleen.

**Figure 7.**
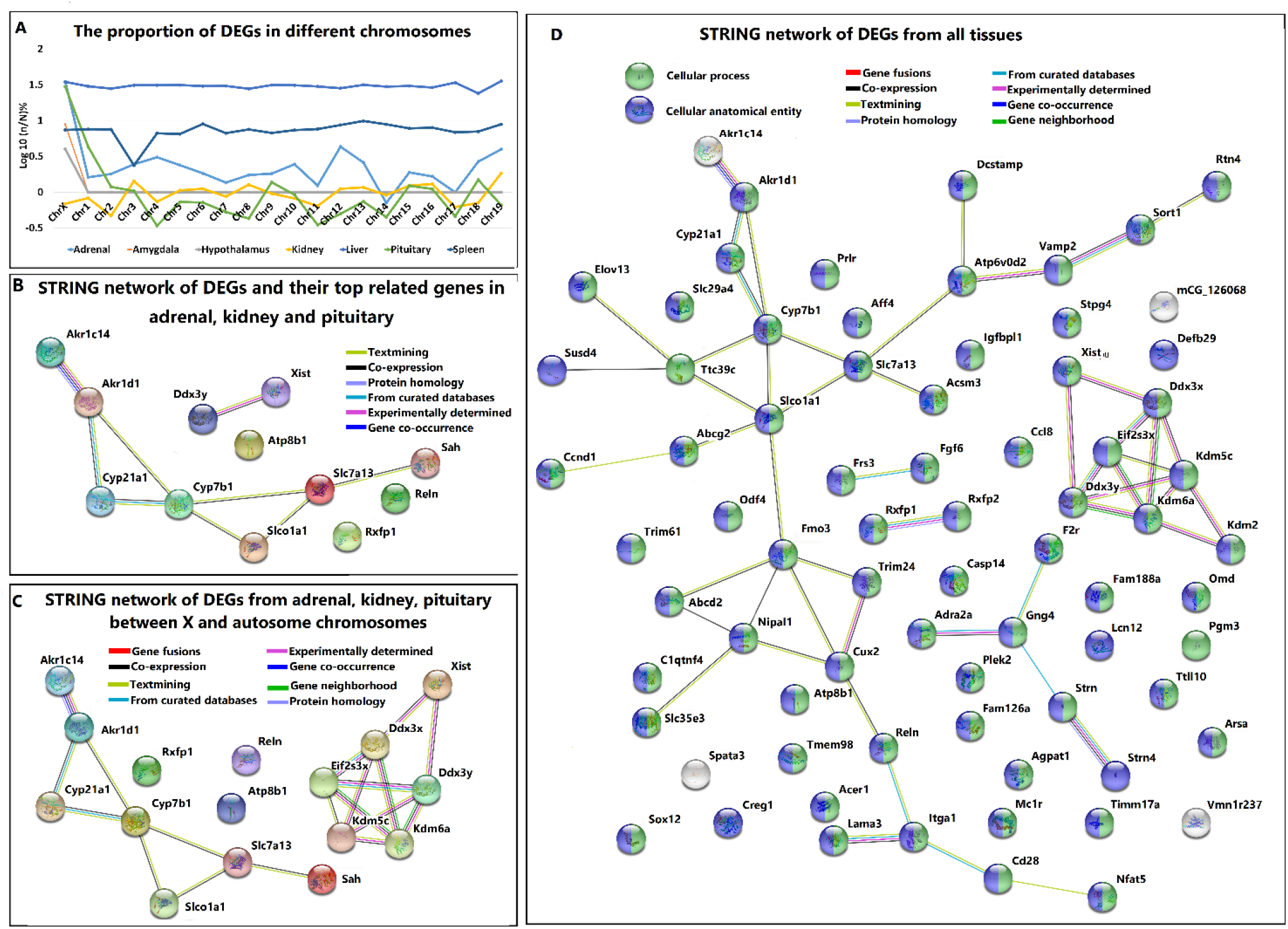
DEGs in mouse autosomal and X chromosomes. **A.** The proportions of DEGs of chromosomes in all tissues examined. The x-axis represents mouse autosomal chromosomes 1-19 and the mouse X chromosome, the y-axis shows the proportions of DEGs that were modified by log10, and the colored lines represent the tissues that were examined. **B.** The STRING networks of DEGs on autosomal chromosomes in adrenal gland, kidney, and pituitary gland and their top related genes. The colored nodes represent query proteins and the first shell of interactors and the edges represent protein-protein associations. The DEGs were associated with multiple functions. **C.** The STRING networks of DEGs on mouse autosomal chromosomes or the X chromosome in adrenal gland, kidney, and pituitary gland. The colored nodes represent query proteins and first shell of interactors and the edges represent protein-protein associations. *Ddx3y* and *Xist* were the top related genes for the autosomal chromosomes and were also closely connected with X chromosome DEGs. **D.** The STRING networks of DEGs on mouse autosomal chromosomes or the X chromosome in liver, adrenal gland, kidney, and pituitary gland. The green nodes represent cellular processes, the blue nodes represent cellular anatomical entities, and the edges represent protein-protein associations. The DEGs on autosome chromosomes in liver were connected with those in other tissues.

#### 3.3.2 The DEGs of adrenal gland, kidney and pituitary gland were predicted to exhibit similar functions

With the exception of the liver, we identified only seven DEGs from human autosomal chromosomes and studied these in detail. When we used STRING to analyze the network among these DEGs and their top related genes (Figure 7B), we observed that they have close interactions related to the pathways involving steroid hormone metabolism.

#### 3.3.3 The *Xist* and *Ddx3y* genes may serve to connect the DEGs from autosome chromosomes with those from X chromosomes

When searching for networks between DEGs in autosomal or X chromosomes of adrenal glands, kidneys, and pituitary glands, we found no direct connections between autosomes and X chromosomes. However, the top related genes of *Slc7a13*, *Slco1a1* and *Acsm3*, namely *Xist* and *Ddx3y* had close interactions with DEGs from X chromosomes, suggesting an indirect interaction of DEGs from autosomal and X chromosomes (Figures 2C and 7C).

### 3.4 Human DEGs exhibited expression patterns that were similar to those in mice

#### 3.4.1 Fewer genes in human autosomal chromosomes exhibited sex bias than in mice and the expression trends between males and females were different

***Akr1d1*** expression exhibited sex bias in mouse adrenal gland, kidney, and liver but not in the corresponding human tissues (Figure 8A). However, the expression value of *Akr1d1* in women was higher than that in men and the trend in humans was the same as that observed in mouse (Figures 1A and 8A).

**Figure 8.**
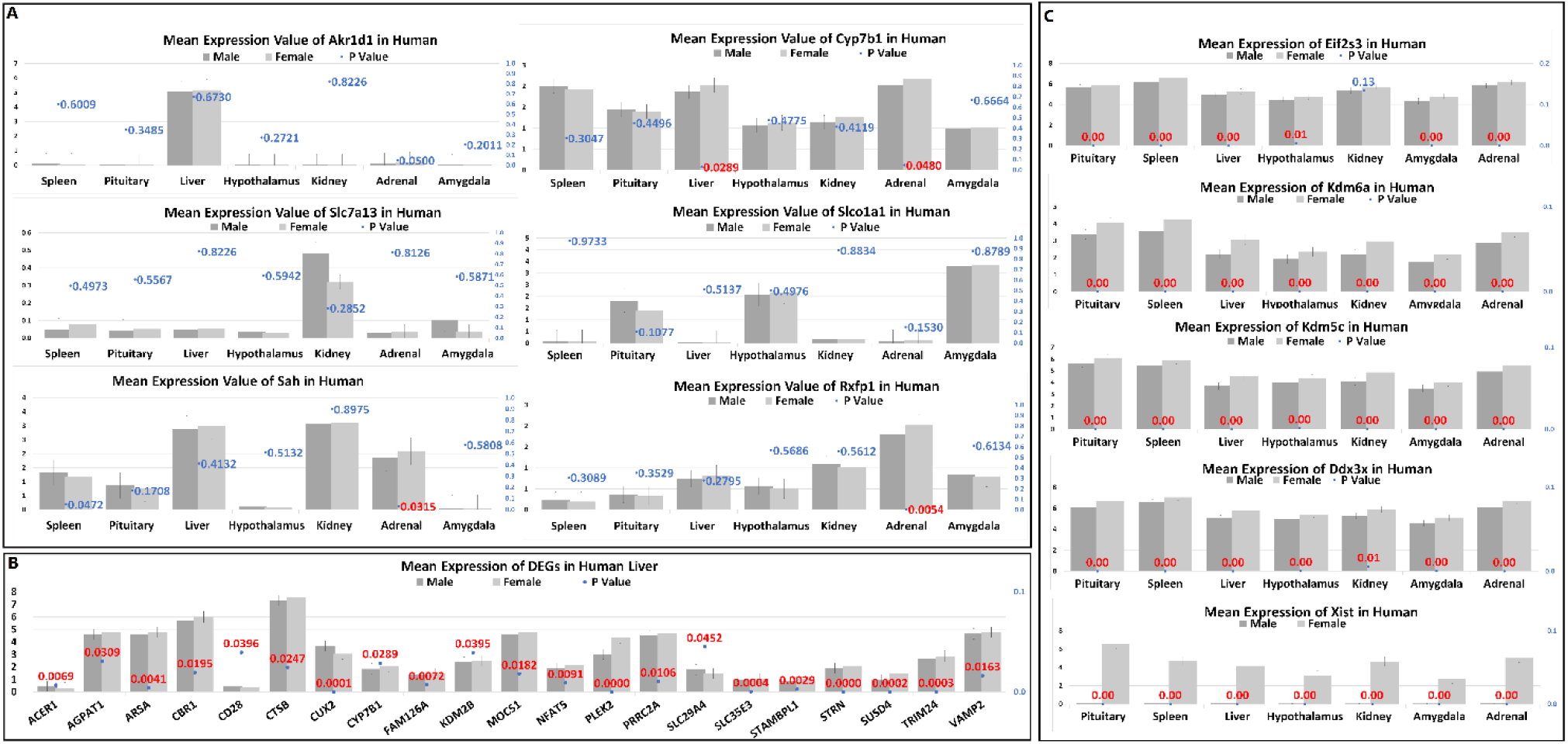
Expression in human tissues of DEGs identified in mouse. **A.** Mean expression values in human tissue of DEGs differentially expressed in mouse adrenal gland, kidney, and pituitary gland from autosomal chromosomes. The panel is labeled as described in the legends to Figures 1A and 3A, except that DEG expression in male and female mice is represented by dark gray and light gray bars, respectively. *Cyp7b1* exhibited sex bias in human liver and adrenal gland, while *Acsm3* and *Rxfp1* exhibited sex bias in the human adrenal gland. **B.** The mean expression values of DEGs identified in liver from mouse autosomal chromosomes whose expression also exhibited sex bias in human liver. The panel is labeled as described in the legends to Figures 1A, 3A**, and** 8A. All of the sexually dimorphic genes we identified in mice also exhibited statistically significant differences in expression in human tissues. **C.** Mean expression values in human tissue of DEGs differentially expressed from mouse X chromosomes. The panel is labeled as described in the legends to Figures 1A, 3A**, and** 8A. Except for *Eif2s3* in kidney, all sexually dimorphic genes that we identified in mice also exhibited sex bias in human tissues.

***Cyp7b1*** expression exhibited sex bias in mouse kidney, pituitary gland, adrenal gland, and liver but in humans only exhibited sex bias in liver and adrenal gland. The expression values of *Cyp7b1* between the sexes in human kidney, liver, pituitary gland, and adrenal gland were opposite those observed in the same tissues from mice (Figures 2B and 8A).

***Slc7a13*** expression exhibited sex bias in kidney, liver, and adrenal glands of mice but not in human tissue. The expression trend of *Slc7a13* between the sexes in human kidney was the same as that detected in mouse kidney, but its expression trend in liver, pituitary gland, and adrenal gland from humans was opposite that detected in these tissues in mice (Figures 2B and 8A).

***Slco1a1*** expression exhibited sex bias in mouse kidney, liver, and amygdala but not in humans. The expression trend of *Slco1a1* between the sexes in human hypothalamus was the same as that in mouse hypothalamus, but in human amygdala was opposite that in observed in mouse amygdala (Figures 2B and 8A).

***Acsm3*** expression exhibited sex bias in mouse kidney and liver but in human tissue, we observed sex bias only in adrenal glands. The expression trend of *Acsm3* between sexes in human kidney, adrenal glands, and liver was opposite that in the corresponding mouse tissue, but the trend was the same for the pituitary gland (Figures 4A and 8A).

***Rxfp1*** expression exhibited sex bias in mouse pituitary gland, kidney, and adrenal gland but only in human adrenal glands. The expression trend of *Rxfp1* between the sexes in human adrenal gland, hypothalamus, and kidney was the same as that detected in mouse tissue, but in human liver, pituitary gland, and spleen, the expression trend was opposite that in the corresponding mouse tissues (Figures 2B and 8A).

Seventy of the DEGs we detected in mouse liver were also expressed in human liver (**Supplementary table 22**). When we compared the mean expression values of these genes in human liver, we detected sex bias for 21 of them (Figure 8B). Moreover, unlike the expression pattern in mouse liver (Figure 5A), most of the DEGs observed in human liver (17/21; 80%) exhibited higher expression in female than male individuals.

#### 3.4.2 The DEGs of human X chromosomes were similar to those in mice

Except for *Eif2s3* in kidney, all X-linked DEGs we detected in mouse tissue also exhibited sex bias in human (Figure 8C). Moreover, except for expression of *Ddx3x*, *Eif2s3x*, *Kdm6a* in autosomal chromosomes in the kidney, all of the DEGs detected in human autosomal and X chromosomes exhibited higher expression in female than in male individuals (Figures 6A and 9).

**Figure 9.**
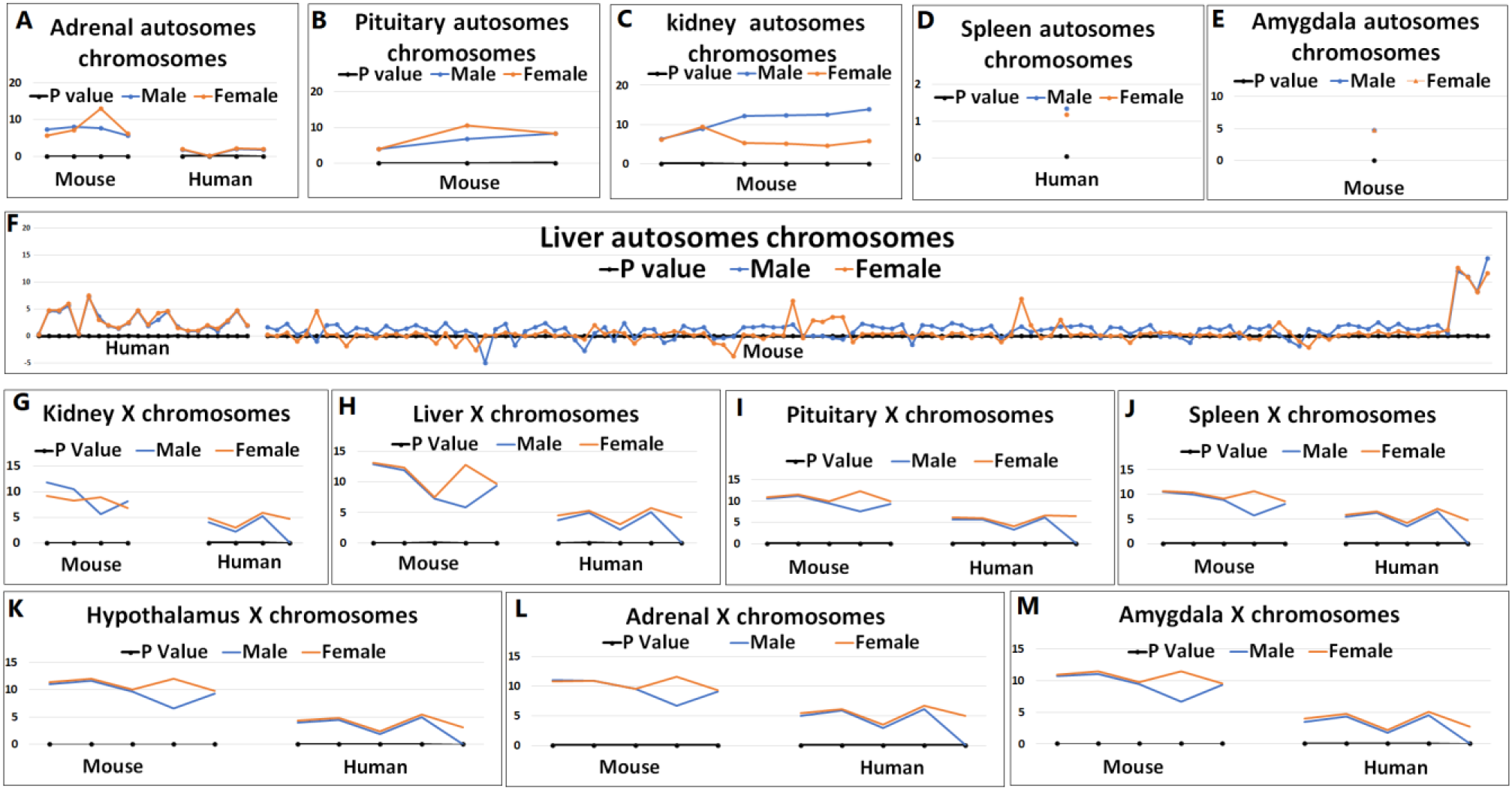
Expression of DEGs in mouse and human autosomal and X chromosomes. A-M, panels show mean expression values of sexually dimorphic genes on autosomal or X chromosomes from the different mouse and human tissues examined. For each panel, mouse data is shown on the left and human data on the right, where appropriate; the x-axis represents the DEGs; the y-axis represents the mean expression value of the DEGs; the blue line represents expression in males; the tangerine line represents expression in females; and the black line represents the p value of differences between expression in males and females from the student’s t-test (*p* values < 0.05 are statistically significant). **A.** Mean DEG expression from mouse and human adrenal gland autosomal chromosomes. **B.** Mean DEG expression from mouse pituitary gland autosomal chromosomes. We detected no DEGs in human pituitary gland autosomes. **C.** Mean DEG expression from mouse kidney autosomal chromosomes. We detected no DEGs in human pituitary gland autosomes. **D.** Mean DEG expression from human spleen autosomal chromosomes. We detected only one DEG in human spleen autosomes and none in mouse. **E.** Mean DEG expression in mouse amygdala autosomal chromosomes. We detected only one DEG in human amygdala autosomes and none in mouse. **F.** Mean DEG expression in mouse and human liver autosomal chromosomes. G-M. Mean DEG expression from mouse and human X chromosomes in the following tissues: **G,** kidney; **H,** liver; **I,** pituitary gland; **J,** spleen; **K,** hypothalamus; **L,** adrenal gland; and **M,** amygdala. In all cases, DEG expression values were higher in human tissues than in the corresponding mouse tissues.

### 3.5. Differences and similarities between expression of autosomal and sex chromosome genes

#### 3.5.1. Distribution of DEGs in each tissue between autosomal and X chromosomes

The relative numbers of DEGs on autosomal and sex chromosomes were as follows: **1)** adrenal glands: autosomal, 67 (0.25%); X-Chr, 284 (23.5%); **2)** amygdala: autosomal, 0; X-Chr, 10 (0.83%); **3)** hypothalamus: autosomal, 0; X-Chr, 10 (0.83%); **4)** kidney: autosomal, 86 (0.20%); X-Chr, 3 (0.21%); **5)** liver: autosomal, 3,648 (20.22%); X-Chr, 198 (28.65%); **6)** pituitary gland: autosomal, 19 (0.07%); X-Chr, 249 (20.63%); and **7)** spleen: autosomal, 26 (0.09%); X-Chr, 13 (1.02%), as shown in Table 1. The amygdala and hypothalamus had no sex-specific DEGs on autosomal chromosomes but each had ten on the X chromosome. The kidney had 86 DEGs on autosomal chromosomes but only three on the X chromosomes, which was the smallest number of DEGs in X chromosomes in any of the tissues we examined. For adrenal gland, spleen, and pituitary gland, more DEGs were found on the X chromosome than on autosomal chromosomes. Finally, the liver had the most DEGs, yet the proportion of DEGs on autosomal and X chromosomes varied the least (Figure 7A).

#### 3.5.2. Functions/pathways of DEGs in each tissue between autosomal and X chromosomes

We used STRING to analyze the functions/pathways of all DEGs and the top ten of each enrichment are listed by FDR rank in Table 4. DEGs on X chromosomes primarily function as regulators of protein domain and GTP binding, while those on autosomal chromosomes mainly serve as regulators of metabolism. DEGs in the adrenal gland were primarily related to oxidoreductase enzymatic activity, and DEGs in the kidney and liver were related to metabolism of drugs, heme, steroid hormones, and retinol, bile secretion, and chemical carcinogenesis.

**Table 4:**
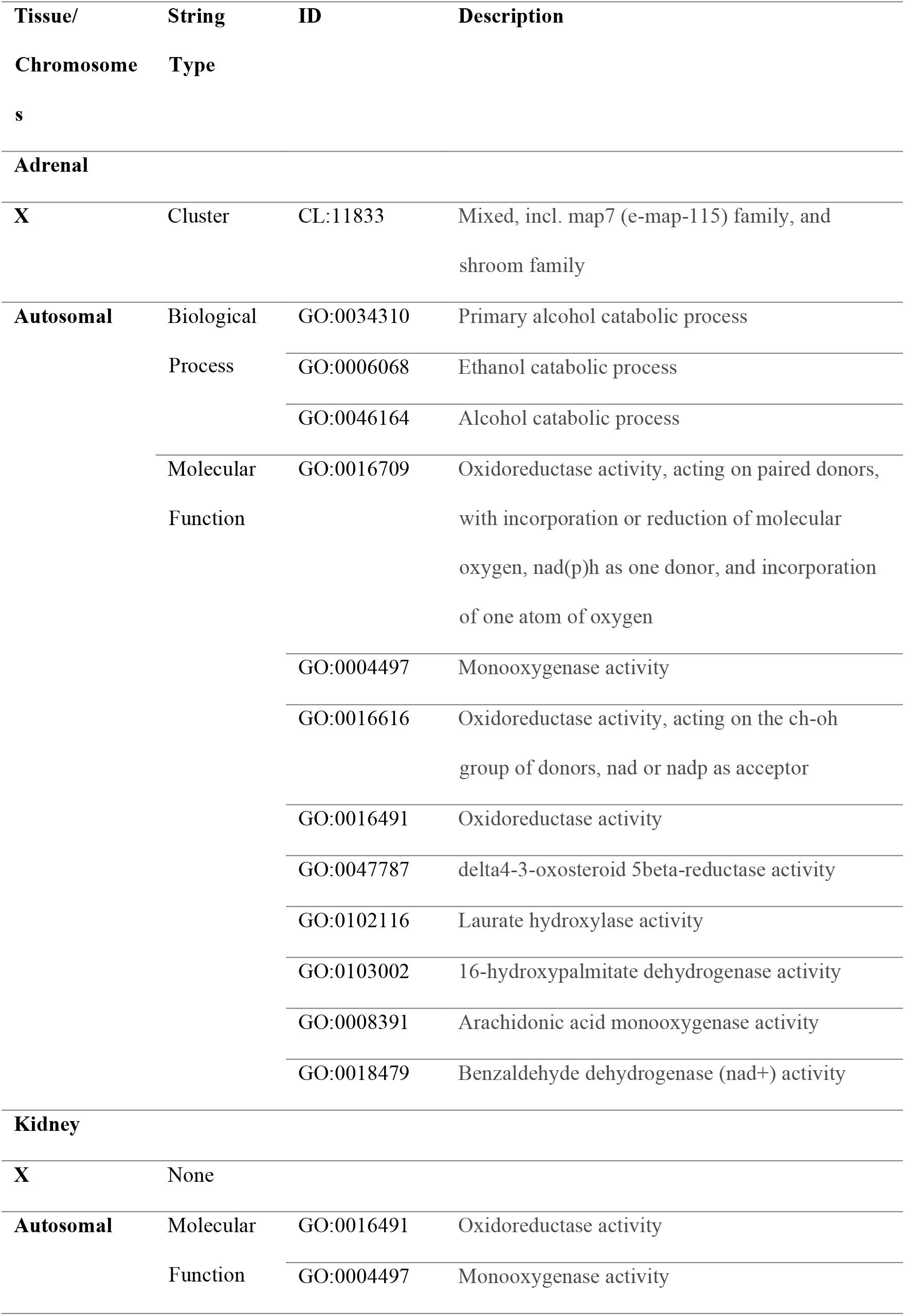
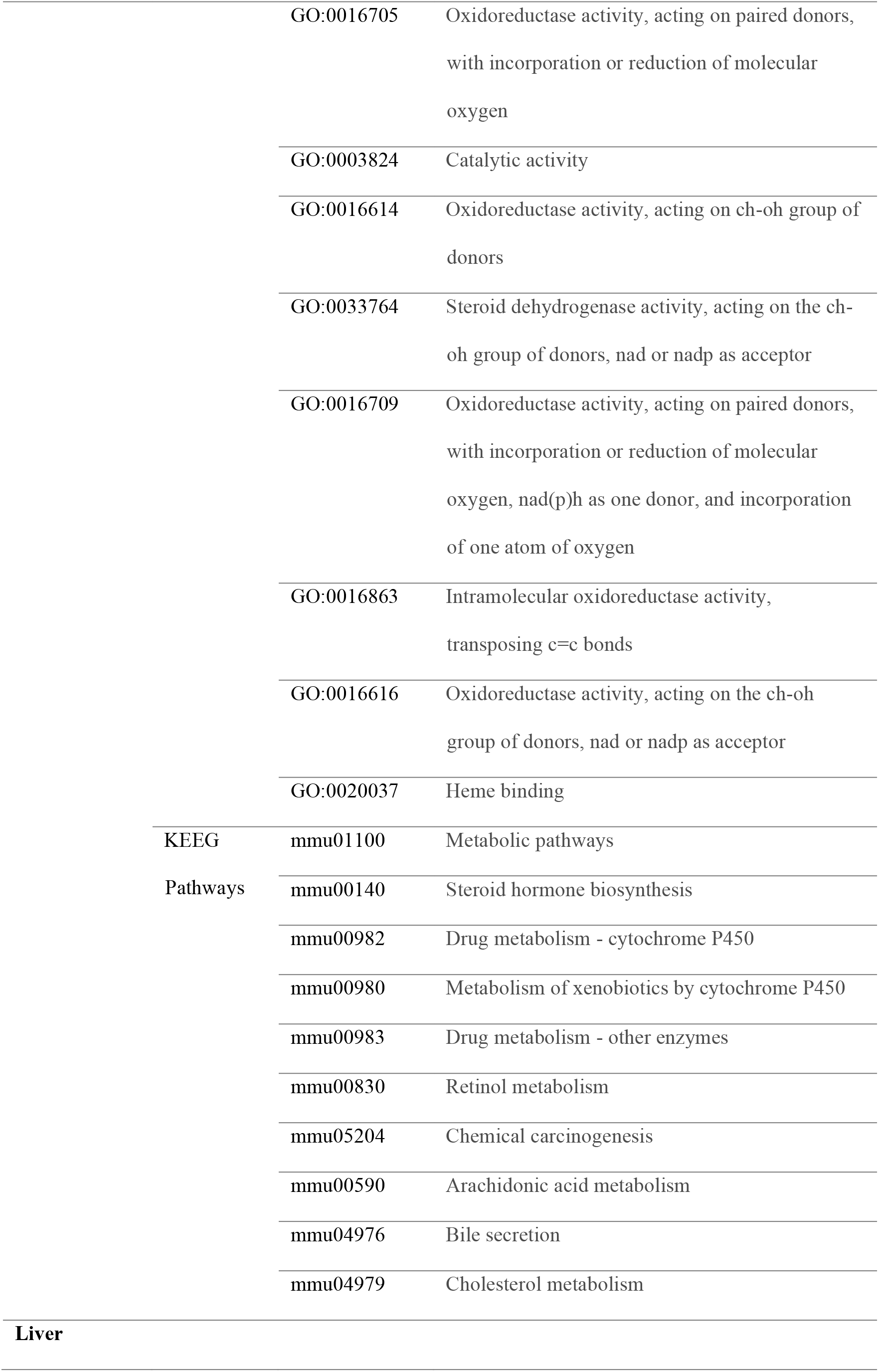
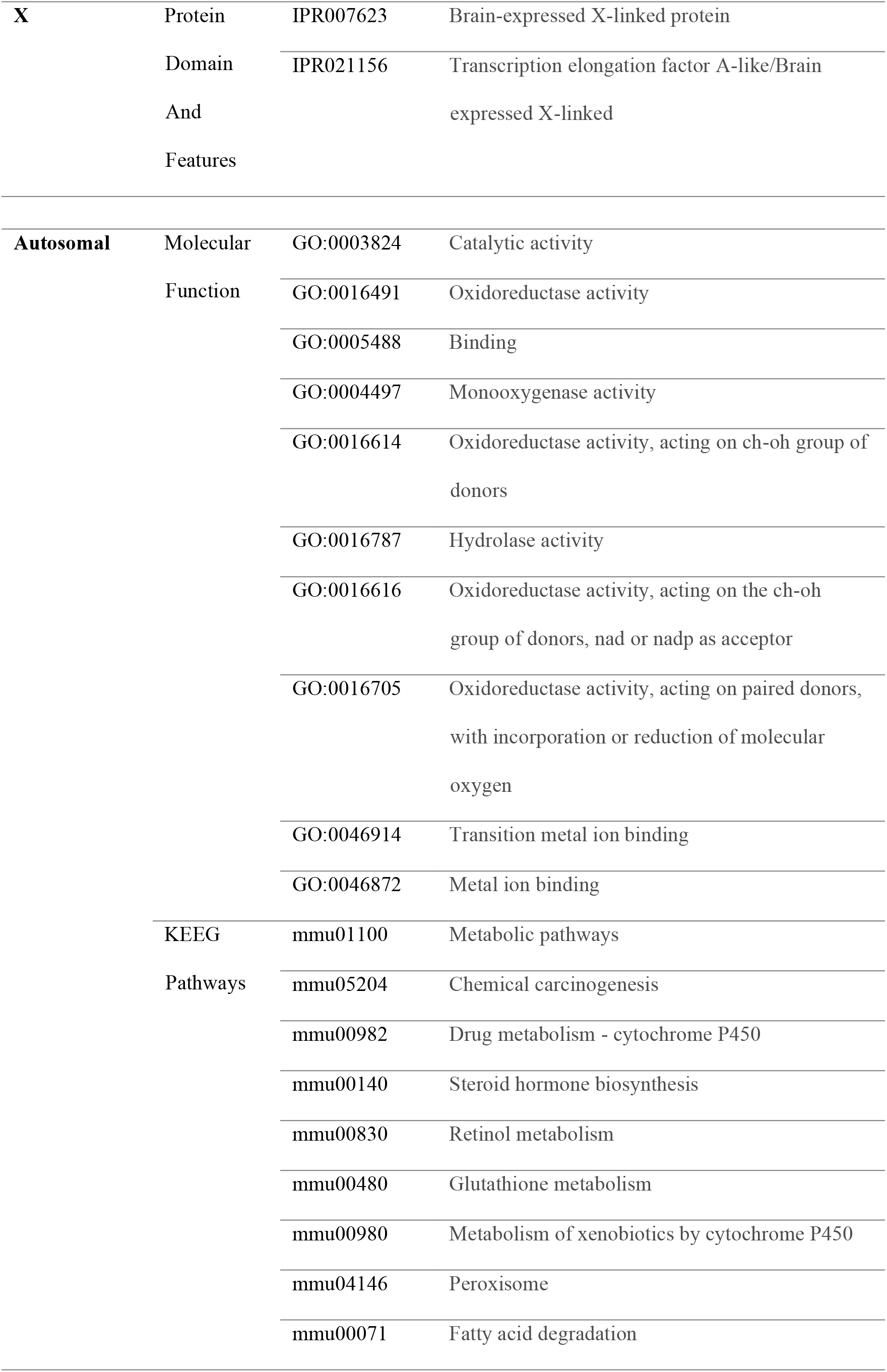
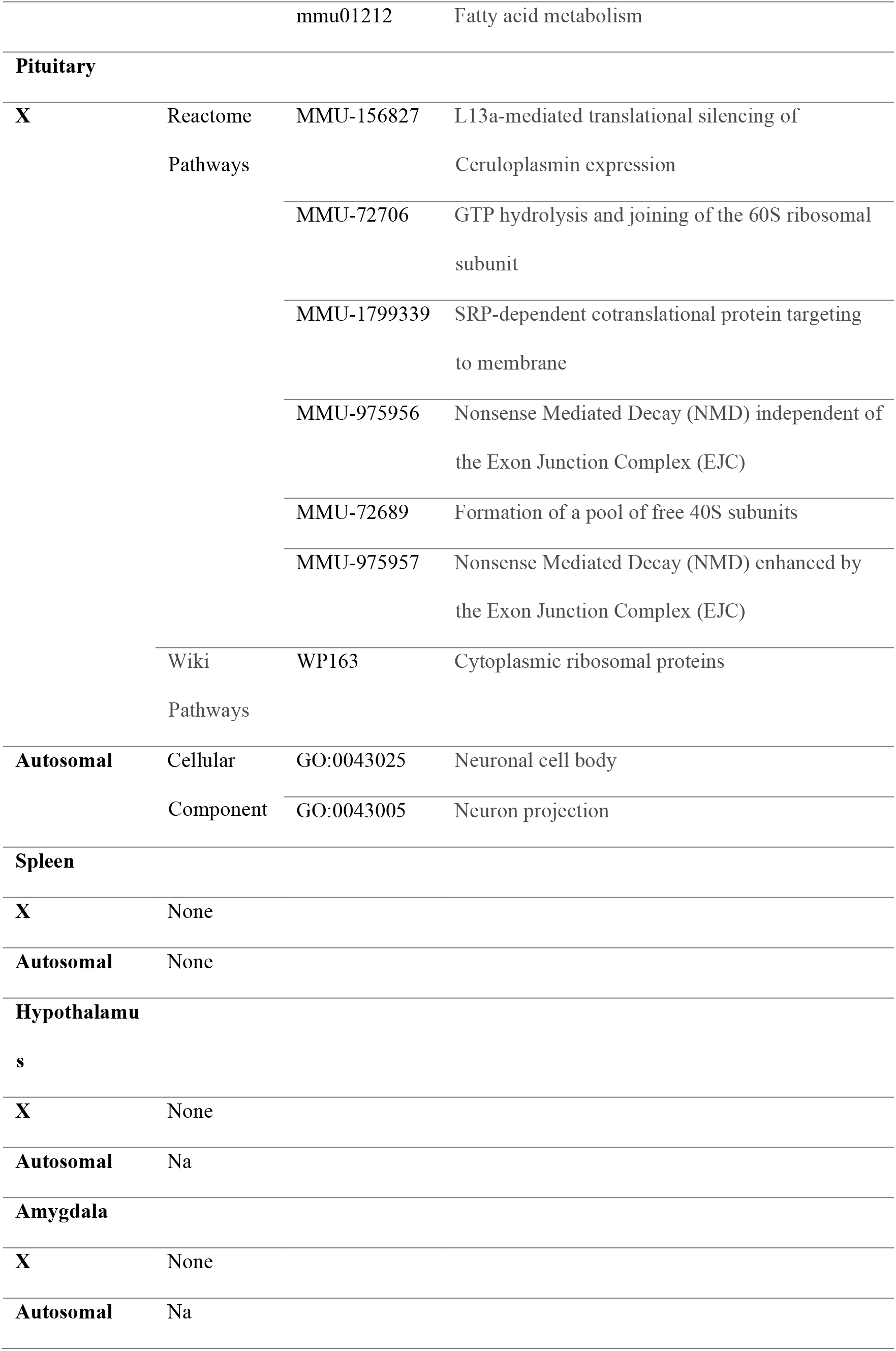
STRING results of sex differential genes between autosomal and sex chromosomes in all tissues.

#### 3.5.3. Structure and function of proteins encoded by DEGs in each tissue between autosomal and X chromosomes

The DEGs with *p* = 0 exhibited sex bias in nearly all tissues we examined (**Supplementary table 28**). Information about the structure and functions of proteins encoded by these genes was extracted from GeneCards (http://www.genecards.org). The following genes and proteins were mapped to the X chromosome: **1)** *Eif2s3x* (GC0XP024054) encodes the largest subunit of a heterotrimeric GTP-binding protein that is involved in the recruitment of methionyl-tRNA(i) to the 40 S ribosomal subunit during translation. **2)** *Kdm6a* (GC0XP044873) encodes a lysine demethylase enzyme that catalyzes the demethylation of tri- or di-methylated histone H3. The protein contains a Jumonji C (JmjC) domain that adopts an evolutionary conserved-barrel structure; such structures often contain binding sites for -ketoglutarate and iron. **3)** *Kdm5c* (GC0XM053176) encodes a protein member of the SMCY homolog family that contains several domains characteristic of transcription factors, including one AT-rich interaction domain (ARID) often involved in DNA binding, a JmjC domain, a Jumonji N (JmjN) domain, and two plant homeodomain (PHD)-type zinc finger domains, often involved in chromatin-mediated regulation of transcription. **4)** The *Xist* gene (GC0XM073820) resides in the X inactivation center on the X chromosome and encodes a noncoding RNA that is involved in inactivation of one of the pair of X chromosomes in females, thereby promoting dosage-dependent equivalence of X chromosome genes between males and females. **5)** *Ddx3x* (GC0XP041333) encodes a protein member of the large DEAD-box family, that is involved in translation, cellular signaling, and viral genome replication. Dysregulation of this gene has been implicated in tumorigenesis.

The important DEGs from autosomal chromosomes (*Akr1d1, Cyp7b1, Sl7a13, Slco1a1, Acsm3,* and *Rxfp1*) and the proteins they encode were described in detail in an earlier section of the paper. These proteins have been related to regulation of steroid hormones, cytochrome P450, members of the G protein-coupled 7-transmembrane receptor superfamily, and synthesis of bile acids and salts.

### 3.6. Difference and similarities among sexually dimorphic gene expression

There are many similarities for the sexual dimorphism of gene expression among different tissues, yet there are more differences between autosomal chromosomes and X chromosomes. Genes on the X chromosome that are expressed in sexually dimorphic manner relate to the early steps of life (developmental genes), while DEGs on autosomal chromosomes are linked to more specific biological functions that mainly focus on control of metabolism pathways.

### 3.7 Candidate regulator genes

From our results it is apparent that: **1)** except for *Masp1*, *Fbxo15* and *Timm21*, all of the candidate genes can regulate the DEGs in the hypothalamus; **2)** six of the ten candidate genes (*Masp1*, *Fbxo15*, *Timm21*, *Mosmo*, *Acsm5*, and *Acsm1*) can regulate the DEGs in the pituitary gland; **3)** an overlapping set of six of the ten candidate genes (*Hao1*, *Plcb4*, *Masp1*, *Mosmo*, *Acsm5*, and *Acsm1*) can regulate DEGs in the liver; **4)** five candidate genes (*Mosmo*, *Acsm5*, *Acsm1*, *Slc12a6*, and *Emc7*) can regulate DEGs in the spleen. The candidate genes in the hypothalamus and pituitary gland may affect the hypothalamic-pituitary axis to regulate the sex-specific DEGs. The liver is the major site of hormone metabolism, which may account for the detection of so many regulator candidate genes in the liver (Langer and Chiandussi, 1987). We identified the only candidate genes for regulation of *Acsm3* in the spleen, namely *Slc12a6*, *Emc7*, *Acsm5*, and *Acsm1*. In light of the observation that the profiles of oxylipin and polyunsaturated fatty acids vary in a sex-specific manner (Pauls et al., 2020), the candidate regulator genes of *Acsm3* may contribute to these sex differences in oxylipin and polyunsaturated fatty acid profiles in spleen.

## 4. Summary and Conclusions

This paper aimed to analyze the difference of sex-specific expression genes across autosome and X chromosome genes in seven tissues using existing data from the BXD family of RI (recombinant inbred) mouse strains. We identified 4.61% of autosomal probes and 15.54% of X chromosome probes whose gene expression values exhibited a sex-associated differential. It is interesting to note that we did not detect a single autosomal gene with differential expression common to the amygdala and hypothalamus, while in liver genes on both autosomal and X chromosomes exhibited significant sexually dimorphic expression. The second highest difference in gene expression across the autosomes was identified in the spleen, where a similar proportion of DEGs were located on the X chromosome. In adrenal gland, liver, and pituitary gland, about 30% genes on the X chromosome exhibited differential expression. In kidney, genes in both autosomal and X chromosomes exhibited a very low proportion of differential expression between sexes. The sexual dimorphism of gene expression exhibited different proportions in different tissues, suggesting that sex-specific differences between humans may influence an individual’s response to human disease to varied levels in different tissues.

Our study is the first to show sexual dimorphism in gene expression in RI mouse strains, where the proportion of genetic variation may approximate that of natural mouse populations. Through our study, we searched through hundreds of genes, most of which have been previously well studied. By performing QTL Interval Mapping, we identified the candidate regulators of DEGs that vary by sex. Incorporating Network Graph and STRING network analysis, we studied the correlations between the DEGs we identified and their candidate regulators in different tissues. The genes *Xist* and *Ddx3y* may serve to connect the DEGs on autosome chromosomes with those on X chromosomes, which suggest that the differential gene expression we studied was associated with sex differences. The candidate DEG regulator genes *Hao1*, *Plcb4*, *Mosmo*, *Acsm1*, and *Emc7* are related to important pathways and human diseases, but it had not yet been determined whether the functions of the proteins they encode were related to sex bias or not.

We also compared the DEGs we identified in mice with those in humans. We found that the expression of DEGs on mouse X chromosomes exhibited similar patterns on human X chromosomes but that the expression trends for genes on mouse autosomal chromosomes were different than for human autosomal chromosomes. The similarities and differences of sex biased gene expression in mice and humans will inform future studies that extrapolate mouse data to conclusions relevant to human health outcomes and public health.

## 5. Methods

### 5.1 Downloading and organization of data

Gene expression data from adrenal gland, amygdala, hypothalamus, kidney, liver, pituitary gland, and spleen tissue in male and female BXD mice were downloaded from the University of Tennessee Health Science Center (UTHSC) GeneNetwork database, version 1, accessed 03/16/2016 (http://artemis-20160316.genenetwork.org/). These data were collated from the following detailed datasets: INIA Adrenal Affy MoGene 1.0 ST (Jun12) RMA Females/Males; INIA Amygdala Affy MoGene 1.0 ST (Nov10) RMA Female/Male; INIA Hypothalamus Affy MoGene 1.0 ST (Nov10) Female/Male; Mouse kidney M430v2 Female/Male (Aug06) RMA; UNC Agilent G4121A Liver LOWESS Stanford (Jan06) Females/Males; INIA Pituitary Affy MoGene 1.0ST (Jun12) RMA Females/Males; and UTHSC Affy MoGene 1.0 ST Spleen (Dec10) RMA Females/Males. The visual basic application (VBA) macro in Excel and a two-tailed t test was used to compare the mean expression levels in male and female tissues and to calculate *p* values for each gene in each tissue. A low *p* value suggests that there is a small chance that such data could have been obtained given the null hypothesis that mean expression levels are equal in males and females. The data for all the genes and tissues were gathered and another VBA macro was used to identify genes whose *p* values were less than 0.00001 in most or all of the tissues. The false discovery rate (FDR) was analyzed for each chromosome, with the FDR test number set equal to the gene number of each chromosome, and a q value (FDR-adjusted *p* value) of less than 0.05 considered to be statistically significant.

A total of 93 strains of BXD RI mice, their parental C57BL/6J (B6) and DBA/2 (D2) mice, and hybrid 1^st^ generation B6D2F1, D2B6F1 mice were used in the experiment. The gene chip experiments were completed in Dr. Robert Williams’ lab (UTHSC). Our research aimed to examine the differential expression of genes present on different chromosomes from tissues of both sexes. We comprehensively analyzed genes from autosomal (Chr1∼Chr19) and X chromosomes in the seven tissues indicated above. To access instructive data, we extracted and merged the records obtained with each probe for every chromosome across all tissues. We concatenated these gene expression intensity datasets by retaining data from a total of 206,523 probes common to all cohorts (See Table 1).

### 5.2 Difference analysis

Within autosome and X chromosome genes in seven kinds of tissues, focus was on significant differences, or sex-specific expression genes. Gene expression level modified by log2 data was downloaded from genenetwork.org and used to calculate values to identify differences in expression level between females and males.

The main calculative work involved comparing the levels of gene expression in male and female mice by subtracting the values obtained in female mice from those in male mice. The number of data entries for each gene varied depending on the chromosome or portion of chromosome selected. The mean and standard deviation values were used to perform two-tailed t-tests to quantify the differences in gene expression between females and males. We calculated t statistics, degrees of freedom, *p* values, and raw differences of these datasets. The *p* values represent the probability that a raw difference of magnitude as great as or greater than that observed could be found by chance given the null hypothesis that the population means are equal between males and females. We focused on the genes whose *p* values were less than 0.05 and found some genes that had extremely low *p* value for some tissue.

### 5.3 Tissue comparison

After the data on the sexual dimorphism of gene expression were obtained, the differential expressed genes among different tissues were compared at three levels. First, we compared the total percentages of genes expressed in a sexually dimorphic manner in different tissues for the 19 autosomal mouse chromosomes. Second, we compared the genes that were expressed in a sexually dimorphic manner on the 19 autosomal mouse chromosomes among the different tissues. Third, we compared genes that were expressed in a sexually dimorphic manner on the mouse X chromosome among different tissues. Similarities are calculated with R values from correlation analysis and *p* values from paired 2-tailed Student’s t tests using an Excel spreadsheet.

To normalize the relative expression levels of genes in different tissues, we utilized the actin protein as the standard control in all the tissues. The relative expression level of each gene in a tissue was expressed as the ratio of its expression to the level of actin expression in the tissue.

### 5.4 Chromosome comparison

The genes we identified as being expressed in a sexually dimorphic manner were compared among different chromosomes in three ways. First, we compared the genes on autosomal chromosomes that were expressed in a sexually dimorphic manner in seven tissues with those on the X chromosomes. Second, we compared the genes on autosomal chromosomes expressed in a sexually dimorphic manner from five of the tissues (the amygdala and hypothalamus had no DEGs on autosomal chromosomes). Third, we compared the genes whose locations mapped to different positions of the chromosomes that were expressed in a sexually dimorphic manner. The statistical analysis was the same as that described for the tissue comparisons, above.

### 5.5 Functional analysis of differentially expressed genes

For each key gene, the following procedures were used to analyze its sex differential expression, functions and pathways. **1)** The mean expression levels in both sexes were obtained from each data set of seven tissues from GeneNetwork. The expression patterns of each gene in all strains of both sexes were collected and compared. Data obtained from these analyses include the standard errors of each data set and p values of expression levels between sexes in each tissue. **2)** Matrix comparisons were done using the Matrix function at GeneNetwork. The Gene network of the key gene for all seven tissues in both sexes were constructed with the Graph function. Pearson’s correlation coefficients were calculated by the Network Graph and line threshold show c. Different types (thickness and color) of line represent different kinds of correlation. **3)** Sample r correlations by Pearson rank were used to obtain two additional types of correlations. For each sex, the top 100 most closely correlated phenotypes were obtained and the most correlated phenotype to the gene of each sex was compared to the gene of the other sex. The same analysis was performed with the top 100 most closely correlated genes in the correspondent tissue. **4)** The function of each gene that was expressed in a sexually dimorphic manner and its top 100 most correlated phenotypes were compared to investigate the portion of phenotypic matches to its known function(s). The phenotype(s) and the name of the differentially expressed gene were used as key words to search the PUBMED literature database. The matched publications were carefully read by at least two investigators and evaluated. **5)** The function of each gene that was expressed in a sexually dimorphic manner and its top 100 most correlated genes were compared to investigate the portion of gene matches to its known function. The name of each top gene and that of the differentially expressed gene were used as key words to search PUBMED. The articles obtained in this search were each read by two investigators and analyzed. **6)** The expression QTL (e-QTL) was analyzed for each sex in each of the seven tissues. For the e-QTL at or near significant level, the e-QTL region was further narrowed down to obtain the list of genes that are found within the e-QTL region. **7)** Interval Analyst and Haplotype Analyst functions were used to obtain SNPs and candidate genes that map to the location of chromosomes whose LRS were higher than significance level. The method of identifying candidate genes has been described before (Wang et al., 2014). The details about the SNPs were obtained from Ensembl (http://useast.ensembl.org/) (Cunningham et al., 2022) and details about the candidate genes were obtained from a series of online resources, including GeneCards (www.genecards.org)(Safran et al., 2021) and UniProt (www.uniprot.org) (The UniProt Consortium, 2021). The protein-protein interaction networks functional enrichment analysis of candidate genes list were studied using STRING (https://string-db.org/) (Szklarczyk et al., 2015). **8)** The database for annotation, visualization, and integrated discovery (DAVID, version 6.8, https://david.ncifcrf.gov/) (Sherman et al., 2022) was used for the list of liver DEGs by Gene Ontology (GO) and the Kyoto Encyclopedia of Genes and Genomes (KEGG) pathway analyses. Only enriched gene counts of GO and KEGG terms more than or equal to 3 and p value <0.05 were considered to be statistically significant.

### 5.6. Comparison of amino acid composition

Amino acid sequences of proteins encoded by the genes of interest were uploaded in FASTA format from National Library of Medicine protein database (https://www.ncbi.nlm.nih.gov/protein) (NCBI Resource Coordinators, 2016). FASTA files were uploaded and subjected to amino acid sequence alignment procedure using a built-in function of Molecular Operating Environment (MOE) software (Chemical Computing Group, Montreal, Quebec, Canada). Identity and similarity metrics were used to visualize amino acid sequence comparisons using the protein similarity monitor in the MOE software package.

## Acknowledgments

The authors thank Dr. Robert Williams for the resources at GeneNetwork and for technical assistance with the data analysis. We thank the following people and their colleagues who contributed the data to GeneNetwork: Lu Lu, Kristin M Hamre, Gerald I Byrne, Melloni Cook, Yue Huang, Yueying Zhang, Wenli Lu, Xiaoyun Liu, and Yonghui Ma (UTHSC); Glenn D. Rosen at Beth Israel Deaconess Medical Center; Eric S. Orwoll and Robert F. Klein (Oregon Health and Sciences University); Patrizia Porcu (University of North Carolina School of Medicine); Richard J Webby (St. Jude Children’s Research Hospital); Elissa J Chesler (Oak Ridge National Laboratory/The Jackson laboratory); Gary A. Churchill (The Jackson Laboratory); David Threadgill (Texas A&M University); Arimantas Lionikas (University of Aberdeen); Dan Goldowitz (University of British Columbia); Sara R Jones (Wake Forest University Health Sciences); Richard W. Moyer (University of Florida College of Medicine); Johan Auwerx (School of Life Sciences, École Polytechnique Fédérale de Lausanne); Leslie C. Jones (The Pennsylvania State University); Martina Buresova (Institute of Clinical and Experimental Medicine, Prague); Boris Tabakoff (University of Colorado); Alan D Attie (University of Wisconsin); and Amanda J. Myers (University of Miami).

## Author Contributions

Conceived and designed the experiments: WG, LW, JJ, GW. Performed the experiments: JM, LW, GY, CF, LY, WD, YJ. Analyzed the data: JM, LW, GY, LY, CT, DZL, YJ, HF, CF. Contributed reagents/materials/analysis tools: LL, AB, MJ, WG, GW. Wrote the paper: JM, LW, GY, WG. Revised the manuscript: AB, MJ, HF, LY, WD, DZL, CT, LL, JJ, YJ, GW, WG. Read and approved the manuscript: All authors.

